# Coenzyme A governs proinflammatory macrophage metabolism

**DOI:** 10.1101/2022.08.30.505732

**Authors:** Greg. A. Timblin, Kevin. M. Tharp, Johanna ten Hoeve, Daniel S. Kantner, Ilayda Baydemir, Eric A. Noel, Chandra Khantwal, Pankaj K. Singh, Joshua N. Farahzad, Jorge Domínguez-Andrés, Russell E. Vance, Nathaniel W. Snyder, Valerie M. Weaver

## Abstract

Toll-like receptor (TLR)-dependent macrophage responses rely on acute increases in oxidative mitochondrial glucose metabolism that epigenetically support rapid proinflammatory transcriptional programming via histone acetylation. Subsequent suppression of oxidative metabolism restrains this metabolic-epigenetic support of proinflammatory gene transcription to enforce tolerance, an immunosuppressed state of innate immune memory. Identifying biology that promotes or counters these metabolic-epigenetic changes will inform therapeutic approaches to influence proinflammatory, antimicrobial, and immunosuppressed myeloid cellular states. Here, we demonstrate that Coenzyme A (CoA) is a “metabolic adjuvant”, as supplying exogenous CoA to macrophages both enhances the magnitude of TLR-driven proinflammatory and antimicrobial responses, and reverse tolerance, via promotion of oxidative metabolism. Extracellular CoA, which we isotopically trace to show its direct uptake by macrophages, works synergistically with tonic TLR signaling, which we demonstrate is a critical regulator of nutrient uptake, metabolism, histone acetylation, and gene expression in macrophages. Together, TLR signaling and exogenous CoA promote mitochondrial glucose oxidation, acetyl-CoA production, and TLR target gene-specific histone acetylation, enhancing metabolic-epigenetic support of proinflammatory transcriptional programming. Exogenous CoA unlocks tumor-associated macrophage (TAM)-dependent TLR agonist anti-tumor activity in an *in vivo* breast cancer model, and promotes macrophage restriction of the intracellular bacterial pathogen *Legionella pneumophila in vitro* via an *Irg1*-dependent antimicrobial state of CoA-augmented itaconate biosynthesis. Our findings demonstrate direct acquisition of intact extracellular CoA, and the ability of this exogenously supplemented metabolic cofactor to augment a key oxidative metabolic-epigenetic pathway supporting proinflammatory and antimicrobial macrophage phenotypes. This may inform host-targeted metabolic adjuvant therapies to reverse myeloid immunosuppression.

## Main Text

Macrophage Toll-like receptor (TLR)-dependent proinflammatory responses to pattern- and damage-associated molecular patterns (PAMPs, DAMPs) are crucial for host anti-tumor and anti-pathogen immunity, and for initiating inflammation following tissue injury (*1*). Tolerance, a form of innate immune memory whereby macrophages chronically exposed to PAMPs/DAMPs enter an immunosuppressed state and are refractory to TLR restimulation (*2*), both protects the host from excessive inflammation, and activates pro-resolving, wound healing programs necessary for tissue repair at infection/injury sites (*3–8*). However, this immunosuppressed state can work against the host in certain instances (e.g. cancer, sepsis, chronic infection) if the initial threat is not eliminated, or new threats emerge (*9–12*).

Macrophage proinflammatory responses, and subsequent tolerance, to the TLR4 ligand lipopolysaccharide (LPS) have been intensively studied (*13–15*). TLR4 target gene enhancers and promoters possess low level occupancy of the histone acetyltransferase (HAT) p300, histone H3 lysine 27 acetylation (H3K27ac), and occupancy of the general transcriptional machinery including RNA polymerase II (Pol II) (*16–20*). LPS induces instantaneous increases in glucose uptake (*21*) and activation of signal-dependent transcription factors (SDTFs), including NFκB. Following partial oxidation to pyruvate via glycolysis, glucose-derived carbons are further oxidized via the mitochondrial TCA cycle, which requires pyruvate dehydrogenase (PDH)-mediated acetyl-CoA production in the mitochondrial matrix. ATP citrate lyase (ACLY) then cleaves TCA cycle-derived citrate into nucleocytosolic acetyl-coA (*22*, *23*). In parallel, SDTFs bind TLR4 target gene enhancers and promoters, augmenting HAT recruitment, which utilize nucleocytosolic acetyl-CoA to rapidly (i.e. within minutes) increase histone acetylation (*17*, *19*, *22–24*). This facilitates biophysical loosening of histone-DNA contacts, and promotes co-factor recruitment (*19*) that transition these primary response genes (PRGs) to an activate state of high H3K27ac and elongating Pol II. (*18*, *19*, *25*, *26*). Following this oxidative burst, a shutdown of mitochondrial oxidative metabolism occurs. This is a consequence of oxidative damage and inhibition of PDH, TCA cycle enzymes, and the electron transport chain (ETC), caused by reactive molecules produced during the oxidative burst (*27–33*). This decreased mitochondrial oxidative metabolism enforces tolerance via suppressing the glucose-derived acetyl-CoA production that initially supported histone acetylation and proinflammatory gene activation (*22*, *34–36*). Concurrently, aerobic glycolysis-derived lactate promotes a transition to an anti-inflammatory macrophage phenotype (*6*, *7*, *37*).

As immunosuppressed patient myeloid cells display bioenergetic defects consistent with impaired glucose oxidation (*38*), therapeutic strategies to counter tolerance-enforcing immunometabolic changes merit identification. Coenzyme A (CoA) is a Vitamin B5-derived (pantothenate) coenzyme that is a critical regulator of mitochondrial glucose oxidation and acetyl-CoA production from glucose. Here, we describe the ability of intact, exogenously supplemented CoA to be directly taken up by macrophages from their extracellular environment to augment these processes. Supplemented CoA synergizes with TLR signaling to enhance the magnitude of both basal and TLR4-driven proinflammatory gene expression in naïve macrophages, and reverse immunosuppression in LPS-tolerized macrophages, by facilitating metabolic-epigenetic support of transcription via histone acetylation. This “metabolic adjuvant” activity of CoA unlocks the anti-cancer activity of a TLR4 agonist *in vivo* in breast cancer model, and supports an anti-microbial macrophage state that restricts *Legionella pneumophila* growth in an *in vitro* infection model. Our findings demonstrate that a key metabolic-epigenetic axis supporting macrophage proinflammatory transcriptional programming can be directly augmented via CoA supplementation. Moreover, they provide a potential therapeutic strategy for reversing tolerance via restoring the metabolic-epigenetic support of proinflammatory gene transcription.

### CoA supplementation enhances basal and TLR-induced proinflammatory gene expression and reverses tolerance *in vitro*

We previously discovered that CoA supplementation restored LPS-dependent induction of proinflammatory genes including *Il1b* and *Nos2* in macrophages treated with various anti-inflammatory small molecules. This included the endogenous anti-inflammatory estrogen metabolite 4-hydroxyestrone (4-OHE1), which acutely decreased intracellular CoA and acetyl-CoA levels, impairing LPS-induce histone acetylation and proinflammatory gene induction (*34*) (Fig. S1A). That exogenously supplied CoA could overcome the pseudo-starved state of low acetyl-CoA caused by 4-OHE1 to rescue proinflammatory gene transcription was of interest, thus we sought to understand the mechanism. Our studies here focus on timepoints consistent with the acute transcriptional programming of macrophages by LPS via TLR4 (0-4h), as these timepoints coincide with and are relevant for studying the initial metabolic and histone acetylation changes that support proinflammatory gene induction. Moreover, they are timepoints when LPS enhances glucose oxidation via the TCA cycle as measured by ^13^C_6_-glucose tracing into TCA cycle intermediates, but preceding the aforementioned increases in steady-state levels of these TCA intermediates (such as citrate and succinate) that signify a damaged TCA cycle in which these metabolites are consumed at lower rates and accumulate (Fig. 1A).

**Figure 1.**
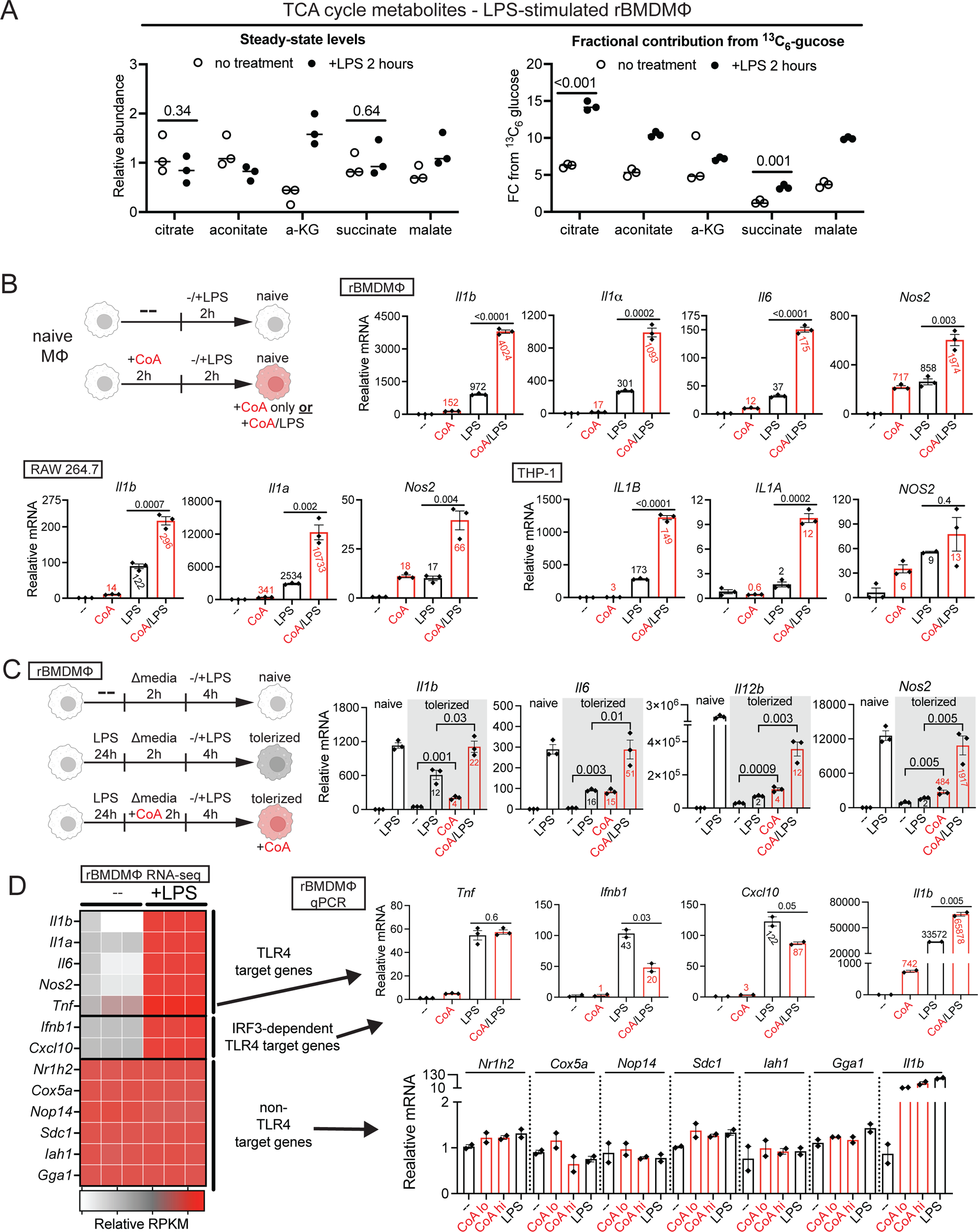
CoA reverses tolerance and enhances TLR-dependent proinflammatory responses *in vitro*. **A.** Steady-state levels of (left) and relative ^13^C_6_-glucose fractional contribution (FC) to (right) select TCA cycle metabolites in recombinant murine macrophage colony stimulating factor (rMCSF)-differentiated bone morrow-derived macrophages (rBMDMs) stimulated with LPS for 2h. n=3 biological replicates per condition, numbers above bars are Student’s t-test P values (planned comparisons)**. B.** Quantitative realtime PCR (qPCR) for LPS-induced proinflammatory gene expression in naïve rBMDMs, RAW 264.7s, and THP-1s with CoA supplementation (250μM). Numbers denote fold-induction over the untreated condition. **C.** qPCR for LPS-induced proinflammatory gene expression in naïve or tolerized bone morrow-derived macrophages (BMDMs) with CoA supplementation (2.5mM). Numbers denote fold-induction over the tolerized, untreated condition. **D.** left – RNAseq heatmap denoting relative expression (RPKM) of select genes in untreated or LPS-stimulated (6h) rBMDMs from ref. 34. top right - qPCR for LPS-induced proinflammatory gene expression with CoA supplementation as described in **B**. bottom – qPCR for gene expression in cells supplemented with 500μM (lo) or 2.5mM (high) CoA for 2h. For all qPCRs, n=2 or 3 biological replicates per condition, mean ± s.e.m is shown, numbers above bars are Student’s t-test P values (planned comparisons).

In naïve primary bone marrow-derived macrophages differentiated with recombinant murine macrophage colony stimulating factor (MCSF) in DMEM media (referred to as rBMDMs), CoA supplementation alone (250-500μM CoA, in line with average intracellular concentrations (*39*)) was sufficient to significantly upregulate transcript levels of the proinflammatory cytokines *Il1b*, *Il1a*, and *Il6*, as well as *Nos2* (Fig. 1B). Furthermore, CoA supplementation strongly synergized with LPS to enhance proinflammatory transcriptional output, displaying non-additive effects when combined with TLR4 stimulation (Fig. 1B). These proinflammatory effects of CoA alone, and CoA synergy with LPS, were also observed in RAW 264.7 murine macrophages, human THP-1 monocytic cells (Fig. 1B), and in BMDMs differentiated with 3T3-L1 cell supernatant in RPMI media (referred to as supBMDMs, Fig. S1B). Immunostimulatory effects of CoA also extended to naïve primary human myeloid cells, as CoA dose-dependently enhanced *IL1B*, *IL1A*, *IL6*, and *TNF* expression in donor monocytes either alone, or in combination with LPS (Fig. S1C). Importantly, quantification of endotoxin levels provide evidence the proinflammatory effects of CoA are not due to contamination of the yeast-derived CoA preparations (Fig. S1D). Back calculations (see methods) estimate that in treating macrophages with 500μM of CoA, they are exposed to approximately 0.1-1 pg/mL of contaminating endotoxin depending on the CoA source (Fig.S1D). As this is 50-500-fold lower than LPS concentrations that produce a quantifiable increase in proinflammatory gene transcription in murine BMDMs (*40*), the robust upregulation of proinflammatory gene transcription in response to 500μM CoA we observed for CoA from all vendors is via a mechanism other than TLR4 agonism by contaminating endotoxin. Moreover, concurrent co-treatment of CoA and LPS significantly and synergistically enhanced the LPS response (Fig. S1E.). This demonstrates the augmentation of the LPS response by CoA is not a consequence of previously described “priming”, whereby ultra-low concentrations of LPS do not themselves elicit measurable responses, but do prime macrophages for enhanced LPS responses in a manner that requires several hours of pretreatment (*40*).

Given its potent immunostimulatory effects, we hypothesized that CoA might reverse LPS tolerance. Consistently, CoA supplementation alone (2.5mM, in line with intramitochondrial concentrations (*39*)) was sufficient to elevate proinflammatory transcript levels in tolerized rBMDMs (Fig. 1C). Moreover, CoA synergized with LPS to partially or completely restore induction of genes encoding proinflammatory genes in tolerized rBMDMs to levels observed in naïve rBMDMs treated with LPS (Fig. 1C). However, *Tnf* tolerance was not rescued by CoA (Fig. S1F).

Further interrogation of the effects of CoA supplementation in naïve rBMDMs suggested that CoA modulates not all, but rather specific aspects, of TLR4-induced gene expression (Fig. 1D). *Tnf* is unique from LPS-induced PRGs/IEGs like *Il1b* in that its CpG island-rich promoter affords strong inducibility independent of chromatin remodeling activities promoted by histone acetylation (*18*). This, along with observations that *Tnf* induction in murine BMDMs is insensitive to inhibitors or genetic perturbations that prevent mitochondrial glucose oxidation and suppress *Il1b* induction (*32*, *41–43*), suggests *Tnf* might be less dependent on glucose-derived, acetyl-CoA-dependent histone acetylation for induction. Unlike *Il1b*, *Tnf* induction by LPS was not enhanced by CoA in rBMDMs (Fig. 1D). Proinflammatory gene induction by LPS involves modular programs to tune the specific nature of the host defense response. TRIF signaling downstream of TLR4 activates a specific subset of proinflammatory genes involved in antiviral defense in a manner dependent on the transcription Interferon Regulatory Factor 3 (IRF3) for maximal induction, including the Type I Interferon-encoding gene *Ifnb1,* and *Cxcl10* (*25*). In contrast to *Il1b*, CoA did not strongly induce *Ifnb1* or *Cxcl10* alone, and actually repressed their induction by LPS, in rBMDMs (Fig.1D).

Finally, while TLR4 agonism enhances expression of signal-responsive genes involved in host defense, the majority of active genes in macrophages are unchanged in their expression post-LPS stimulation. From our existing RNAseq data (*34*), we chose 6 moderately expressed, non-LPS-induced genes at random, and found that, unlike TLR4 target genes such as *Il1b*, CoA had no effect on their basal expression in rBMDMs (Fig. 1D). This result further underscores that the effects of CoA on macrophage gene expression are specific to a subset of TLR4 target genes, and are not a consequence of a general, global increase in transcription across all active genes in macrophages.

### Macrophages directly take up exogenous CoA

Given the importance of mitochondrial glucose oxidation and glucose-derived acetyl-CoA in epigenetic activation of proinflammatory gene transcription (*22*, *23*, *34*), we proposed the following model for the rapid effects we observed on gene expression by CoA (Fig. 2A). In naïve macrophages, low levels of tonic TLR signaling cooperate with endogenously synthesized CoA to support the maintenance of enhancers/promoters and TLR4 target genes in their poised state characterized by basal occupancy of HATs and low levels of histone acetylation and transcription. Upon exogenous CoA supplementation, this CoA is directly taken up by naive macrophages and enhances their basal capacity for mitochondrial glucose oxidation and acetyl-CoA production. In turn, this fuels HAT-dependent histone acetylation and tunes-up basal levels of transcription specifically at a subset of TLR4 target genes. When CoA is supplemented to macrophages that are then treated with LPS, strong TLR signaling synergizes with newly acquired exogenous CoA to activate proinflammatory gene transcription to levels above that which could be achieved by either CoA supplementation or TLR4 agonism alone. This model generates a set of four testable hypotheses (Fig. 2A). The first hypothesis we set out to test was that macrophages can directly take up extracellular CoA.

**Figure 2.**
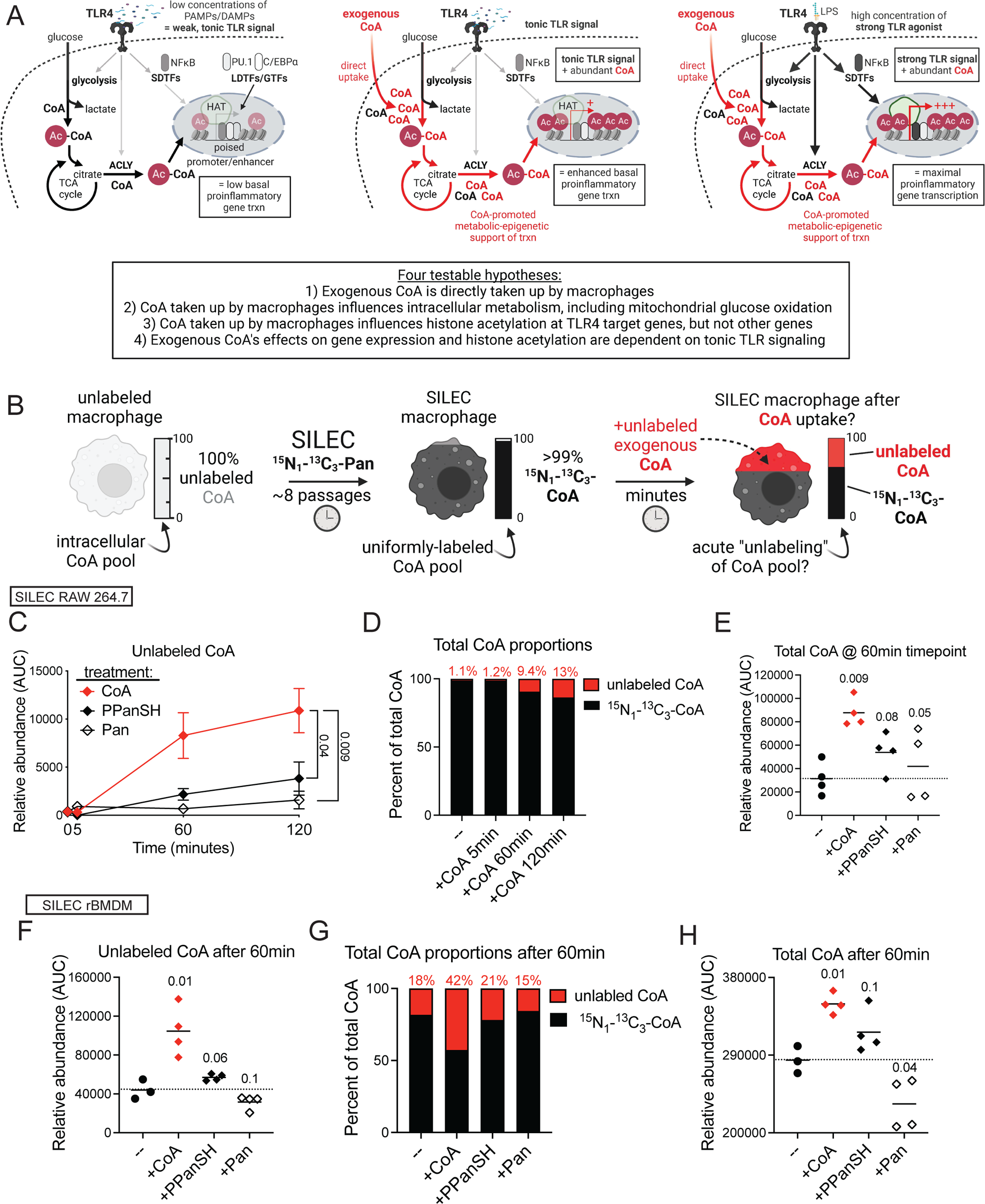
Macrophages directly take up exogenous CoA. **A**. Model for proinflammatory effects of CoA, and testable hypotheses from the model. **B**. SILEC strategy for isotopic labeling of the endogenous CoA pool, and tracing of unlabeled CoA into macrophages. **C**. Abundance of unlabeled CoA over time in SILEC RAW264.7 macrophages treated with unlabeled CoA, PPanSH, or Pan (all 500μM). n=4, mean ± s.e.m. is shown. **D**. For CoA-treated cells from **C**, unlabeled CoA as a percentage of the total CoA pool over time (set to 100%). **E**. For cells from **C**, the total amount of CoA (labeled + unlabeled) at 60 minutes following CoA, PPanSH, or Pan treatment. Mean is shown, numbers above data are Student’s t-test P values (planned comparison with untreated condition). **F**. Abundance of unlabeled CoA in SILEC rBMDMs 60 minutes after treatment with unlabeled CoA, PPanSH, or Pan all 500μM). n=4, mean is shown, numbers above data are Student’s t-test P values (planned comparison with the untreated condition). **G**. For CoA-treated cells from **F**, unlabeled CoA as a percentage of the total CoA pool over time (set to 100%). **H**. For cells from **F**, the total amount of CoA (labeled + unlabeled) at 60 minutes following CoA, PPanSH, or Pan treatment. Mean is shown, numbers above data are Student’s t-test P values (planned comparison with untreated condition).

CoA is thought to be exclusively acquired via intracellular biosynthesis from precursors acquired from the extracellular environment via diet or CoA degradation, namely pantothenate (Pan, Vitamin B5) and 4-phosphopantetheine (PPanSH) (*44*, *45*) (Fig. S2A). Neither Pan or PPanSH supplementation phenocopied CoA’s ability to upregulate basal proinflammatory gene expression in macrophages, or synergize with LPS (Fig. S2B). The CoA degradation products adenosine 3’,5’ diphosphate and dephospho-CoA also lacked these activities (Fig. S2B). Moreover, preincubation of CoA with non-heat inactivated fetal bovine serum containing active ectonucleotide pyrophosphatase/phosphodiesterase (ENPP) enzymes that degrade CoA to PPanSH (*44*) significantly abrogated CoA’s ability to upregulate *Il1b* expression (Fig. S2C), an effect reversed by serum heat inactivation that kills ENPP activity. Thus, no diet-derived or extracellularly-generated CoA degradation product phenocopied CoA’s acute effects of macrophage proinflammatory gene expression.

This prompted us to develop a system to test whether exogenous CoA could be directly taken up by macrophages and significantly contribute to the endogenous CoA pool in a manner with distinct kinetics and magnitudes than precursors that fuel endogenous CoA biosynthesis. As isotopically-labeled CoA is not commercially available, we turned to a method known as SILEC (Stable Isotope Labeling by Essential nutrients in cell Culture), which exploits the essentiality of Pan in eukaryotic cell growth. This technique, in which the unlabeled Pan in culture media is replaced with ^15^N_1_-^13^C_3_-Pan, can achieve near-complete isotopic labeling of the endogenous CoA pool in yeast (*46*, *47*) and mammalian cell lines in culture (*48*). Following SILEC labeling, this would allow treatment of SILEC macrophages with unlabeled exogenous CoA, or unlabeled Pan and PPanSH, and subsequent extraction and liquid chromatography-mass spectrometry (LC/MS) to quantify the contributions of exogenous CoA/Pan/PPanSH to the endogenous CoA pool by quantifying levels of unlabeled intracellular CoA (Fig. 2B).

Following 8 passages in SILEC media, >98% isotopic labeling of the endogenous CoA pool of RAW 264.7 macrophages was achieved (Fig. S2D). Upon treatment of SILEC RAW 264.7 macrophages with unlabeled exogenous CoA, and extensive washing to remove extracellular CoA before extraction (see methods and Fig. S2E), we observed a time-dependent increase in levels of unlabeled intracellular CoA consistent with direct CoA uptake (Fig. S2F). Encouraged, we repeated this experiment, comparing the effects of treatment with equimolar amounts of unlabeled CoA, Pan, or PPanSH. While PPanSH (but not Pan) supplementation drove a time-dependent increase in levels of unlabeled intracellular CoA on the timescale under study, the kinetics of this “unlabeling” of the intracellular CoA pool were distinct from, and significantly lower in magnitude than, CoA supplementation (Fig. 2C). By 60 minutes, unlabeled CoA accounted for between ∼10-30% of the total intracellular CoA pool depending on the experiment (Fig. 2D, Fig. S2F). Interestingly, exogenous CoA treatment significantly increased the total intracellular CoA pool size (i.e. amount of total unlabeled + labeled CoA), an effect not observed upon PPanSH or Pan supplementation (Fig. 2E). This suggests that exogenous unlabeled CoA, in addition to directly contributing to the intracellular CoA pool, also exerts a positive influence on endogenous ^15^N_1_-^13^C_3_-CoA biosynthesis via an unknown mechanism.

Extending these findings to primary murine macrophages, despite being generally post-mitotic after their initial expansion, we were able to achieve >80-90% isotopic labeling of the endogenous CoA pool of rBMDMs via culture for 1 month with 10 media changes (Fig. S2G, Fig. 2G). Treatment of with SILEC rBMDMs with unlabeled exogenous CoA revealed increased unlabeled intracellular CoA, with this effect again distinct in magnitude from the effects of equimolar PPanSH or Pan supplementation (Fig. 2F, Fig. 2G). And as in the SILEC RAW 264.7s, exogenous CoA significantly increased the size of the total intracellular CoA pool in a manner distinct from PPanSH and Pan (Fig. 2H). Together, these results tracing unlabeled CoA into ^15^N ^13^C_3_-Pan SILEC macrophages support the hypothesis that macrophages are capable of direct uptake of exogenous CoA. Thus, while it currently thought that CoA is exclusively obtained via intracellular biosynthesis (*45*), our data provides evidence that macrophages are the first cell type described that are capable of directly acquiring extracellular CoA from their environment.

### CoA taken up by macrophages enhances mitochondrial glucose oxidation and directly supports acetyl-CoA production

We next sought to test the hypothesis that exogenous CoA directly taken up by macrophages positively influences mitochondrial glucose oxidation and acetyl-CoA production (Fig. 2A). We first tested this in the system where we initially discovered the proinflammatory effects of CoA supplementation: the ability of exogenous CoA to overcome the anti-inflammatory effects of 4-OHE1 and rescue LPS-induced proinflammatory gene expression (Fig. S1A). ^13^C_6_-glucose isotope tracing was employed to assess differences in glucose utilization across metabolic pathways in 4-hydroxyestrone 4-OHE1-pretreated, LPS-stimulated rBMDMs without or with CoA supplementation. Following pretreatments and LPS stimulation, ^13^C_6_-glucose was provided to macrophages for the final 30 minutes of a 1.5 hour LPS stimulation (Fig. 3A). Metabolite extraction and LC/MS metabolomics were performed, and the relative fractional contribution (FC) of ^13^C_6_-glucose to approximately 130 different intracellular metabolites were quantified. Interestingly, while CoA supplementation had no discernable effects on the isotopic labeling of glycolytic intermediates between conditions, the ^13^C_6_-glucose FC to nearly all TCA cycle metabolites was significantly enhanced by CoA supplementation (Fig. 3A). Acetyl-CoA supplementation, which we also showed overcomes the immunosuppressive effects of 4-OHE1 (*34*), phenocopied this effect (Fig. 3A), implying CoA is the active moiety of acetyl-CoA. The enhanced ^13^C^6^-glucose labeling was particularly noticeable for acetyl-CoA and citrate (Fig. 3B), providing direct evidence of enhanced glucose oxidation to support biosynthesis of these metabolites by exogenous CoA taken up by macrophages.

**Figure 3.**
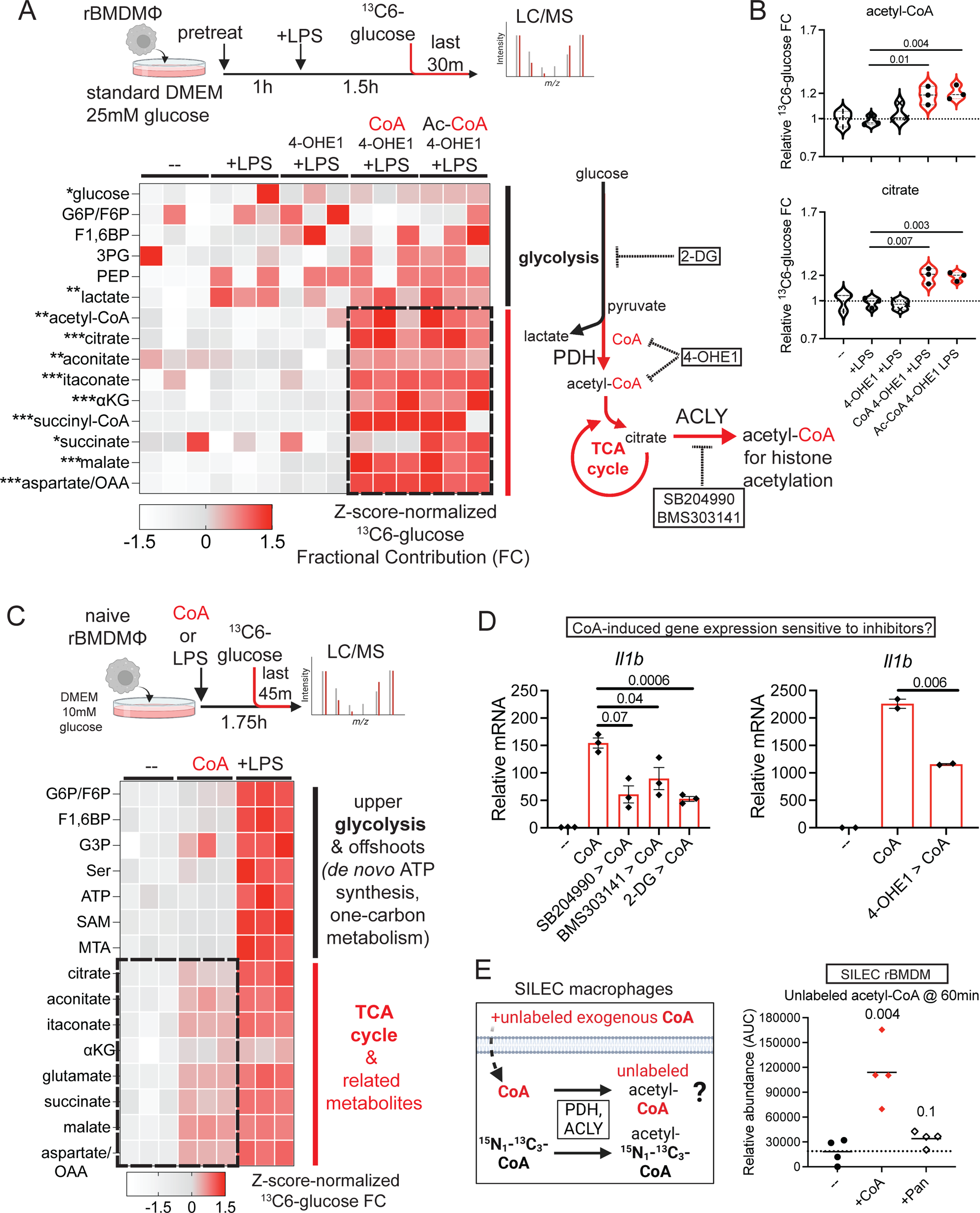
CoA taken up by macrophages enhances mitochondrial glucose oxidation and directly supports acetyl-CoA production. **A**. Top – experimental design for ^13^C_6_-glucose isotope tracing in rBMDMs. Competitive labeling of metabolites with ^13^C_6_-glucose is performed from the final 30 minutes of a 1.5h LPS stimulation to determine how CoA supplementation (250μM) influences glucose labeling of metabolites, serving as a proxy for metabolic pathway activity. Bottom – Heatmap depicting ^13^C_6_-glucose Fractional Contribution (FC) to glycolytic and TCA cycle metabolites, represented as a Z-score for each metabolite. Metabolites with significant FC differences between conditions were identified by One-Way ANOVA. *P<0.05. *P<0.01,***P<0.001. n=3 biological replicates per condition. **B**. Relative ^13^C_6_-glucose FC to acetyl-CoA and citrate from **A**. Numbers above bars are P values from Dunnett’s post-hoc test correcting for multiple comparisons. **C.** Heatmap depicting Z-score-normalized ^13^C_6_-glucose FC to indicated metabolites in naïve BMDMs treated with CoA (250μM) or LPS for 1.75h. **D**. qPCR for CoA-induced *I1lb* expression (2h) in cells pretreated with ACLY inhibitors and 2-DG (left, n=3, mean ± s.e.m), or 4-OHE1 for 1h (right, n=2, mean). Numbers above data are Student’s t-test P values (planned comparison). **E**. left – schematic depicting production of unlabeled acetyl-CoA from unlabeled CoA taken up by SILEC macrophages. right - unlabeled acetyl-CoA levels in SILEC rBMDMs 60 minutes following unlabeled CoA or Pan treatment (both 500μM). n=4, mean is shown, numbers above data are Student’s t-test P values (planned comparison with the untreated condition).

Further supporting the hypothesis that CoA taken up by macrophages enhances mitochondrial glucose oxidation in a second experimental paradigm, CoA alone was sufficient to enhance ^13^C_6_-glucose FC to TCA cycle metabolites in naïve rBMDMs (Fig. 3C). CoA-induced *Il1b* expression in naïve rBMDMs was sensitive to inhibitors that interfere with mitochondrial glucose oxidation, including 2-deoxyglucose, 4-OHE1, and the ACLY inhibitors SB and BMD303141, implying the proinflammatory effects of exogenous CoA are dependent on this CoA’s ability to be taken up and promote mitochondrial glucose oxidation (Fig. 3D). As an orthogonal approach to unequivocally confirm that exogenous CoA is utilized for and supports acetyl-CoA production in naïve macrophages, we tested if unlabeled exogenous CoA provided to naïve SILEC rBMDMs and SILEC RAW 264.7 macrophages could acutely “unlabel” the isotopically heavy endogenous acetyl-^15^N_1_-^13^C_3_-CoA pool in these cells (Fig. S3A). Indeed, unlabeled CoA treatment for 60 minutes resulted in a significant increase in unlabeled acetyl-CoA in both cell types, an effect not observed upon Pan supplementation (Fig. 3E, Fig. S3B). This result definitely demonstrates that exogenous CoA taken up by macrophages is rapidly utilized for endogenous acetyl-CoA biosynthesis.

Finally, we investigated the effects of CoA supplementation in a third experimental paradigm of LPS-tolerized rBMDMs, which accumulate high levels of lactate due to mitoOXPHOS shutdown and a switch to aerobic glycolysis (Fig, S3C). Acute CoA supplementation was sufficient to significantly and rapidly (<4h) reduce steady-state lactate levels (Fig. S3C). Coincident with this effect, CoA drove LPS-tolerized macrophages to use less ^13^C_6_-glucose for lactate production, as indicated by increased in m0 lactate and decreased in m+3 lactate in CoA-supplemented, tolerized rBMDMs (Fig. S3C). These effects are consistent with a model in which a CoA-driven restoration of mitochondrial glucose oxidation in tolerized rBMDMs is responsible for tolerance reversal and the re-establishment of LPS-responsiveness by exogenous CoA.

### Exogenous CoA increases histone acetylation at *cis*-regulatory regions of proinflammatory genes

Having shown that exogenous CoA can be taken up by macrophages to enhance mitochondrial glucose oxidation, we next set out to test the hypothesis that CoA positively influences histone acetylation at TLR4 target genes to increase their expression (Fig. 2A). Prior to strong TLR4 agonism, *cis*-regulatory regions of macrophage proinflammatory genes exist in a state with characteristics of active genes. This includes basal occupancy of transcriptional co-activators such as HATs, low levels of histone acetylation (including H3K27ac), and active transcription. Following strong TLR4 agonism, these genes re “tuned-up” in expression by increased activity of pathways downstream of TLR4, which includes both signal transduction pathways that activate SDTFs (i.e. NFκB) and increase HAT recruitment, and metabolic-epigenetic support via increased glucose-derived acetyl-CoA. The K_m_ for HATs such as p300 lies within physiological acetyl-CoA concentrations (*49*), and increasing p300 activity at specific loci is sufficient to increase both target gene H3K27ac and transcription (*50*). Thus, we hypothesized CoA’s enhancement of mitochondrial glucose oxidation would alone be sufficient to increase basal histone acetylation, and in turn, the basal transcriptional activity, at TLR4 target genes (Fig. 4A).

**Figure 4.**
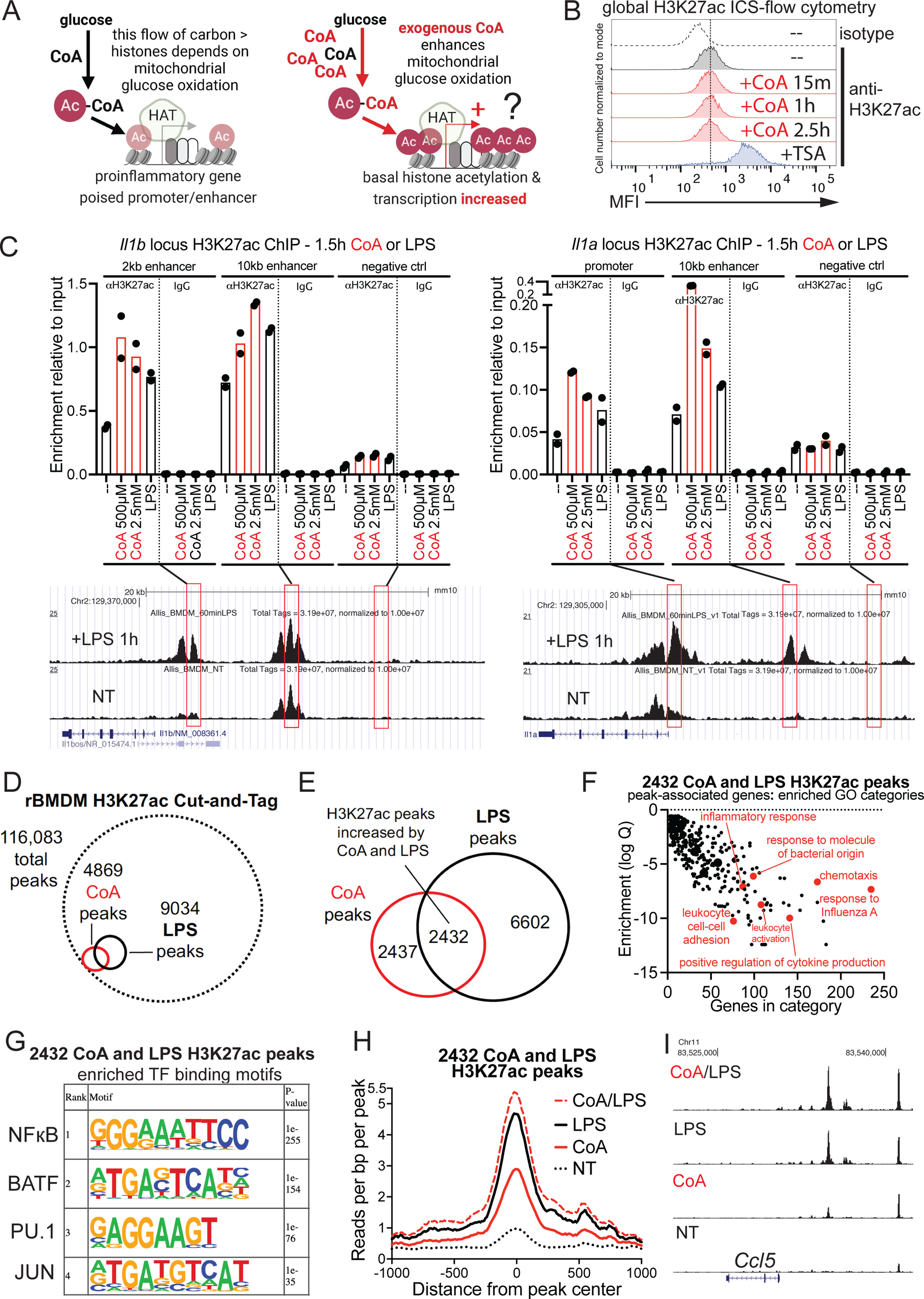
Exogenous CoA increases histone acetylation at *cis*-regulatory regions of proinflammatory genes. **A**. Model depicting how exogenous CoA uptake expands the CoA pool and supports increased acetyl-CoA-dependent histone acetylation at proinflammatory gene promoters/enhancers. **B**. H3K27ac levels measured by intracellular staining (ICS)-flow cytometry in untreated or CoA-treated (500μM) rBMDMs. Treatment with histone deacetylase (HDAC) inhibitor Trichostatin A (TSA, 1μM 18h) serves as a positive control. **C**. H3K27ac levels at *Il1b* and *Il1a cis*-regulatory regions assessed by ChIP-qPCR in control, CoA, or LPS-treated supBMDMs. Mean is shown, each data point represents an independent ChIP experiment/biological replicate (n=2 per condition). IgG control ChIP-qPCR shows specificity of H3K27ac signal. **D**. Venn diagram depicting total number of H3K27ac peaks identified by Cut- and-Tag (CnT) in rBMDMs, relative to the total numbers of peaks increased in read density versus untreated control rBMDMs by either CoA (500μM) or LPS treatment. **E**. Overlap between CoA- and LPS-induced H3K27ac CnT peaks. **F**. Gene ontology (GO) analysis of 2432 genes near CoA- and LPS-induced H3K27ac CnT peaks. **G**. HOMER *de novo* motif finding results for enriched transcription factor motifs in CoA- and LPS-induced H3K27ac CnT peaks. **H**. Cumulative read density at the 2432 H3K27ac peaks in cells treated with either CoA or LPS alone, or CoA/LPS co-treatment. **I**. Response of putative H3K27ac-marked *cis*-regulatory regions in the *Ccl5* locus to treatment with CoA, LPS, or CoA/LPS co-treatment.

H3K27ac, a histone acetylation mark that is directly involved in co-activator recruitment and reports transcriptional activity of genes (*13*, *51*, *52*), is present at low levels in TLR4 target gene promoter/enhancers in resting macrophages, and is increased rapidly upon TLR4 agonism. Consistent with notion that CoA does not nonspecifically and globally increase transcription of all active genes in macrophages (Fig. 1D), CoA supplementation had no effect on global H3K27ac in rBMDMs (Fig. 4B, Fig. S4A). Thus, we turned to targeted techniques to assess CoA’s effects on histone acetylation at specific proinflammatory gene *cis*-regulatory elements. We first used chromatin immunoprecipitation (ChIP) to assess whether CoA alone was sufficient to increase histone acetylation in the *Il1b* and *Il1a* alpha loci, as CoA alone increases the basal transcription of both genes (Fig. 1B). Existing H3K27ac ChIP-seq data (*53*) informed targeted qPCR analysis of regions in these loci that show rapid (<1h) increases in H3K27ac following TLR4 agonism that coincide with increased transcription (Fig. 4C). Indeed, acute CoA treatment alone were sufficient to increase H3K27ac at *Il1b* and *Il1a* promoters and enhancers, but not control regions in these loci (Fig. 4C). This was consistent with our hypothesis, and somewhat surprising since the magnitude of this H3K27ac increase was similar to that observed for LPS treatment, which causes a much stronger increase in *Il1b* and *Il1a* mRNA expression than CoA (Fig. 1B). This is likely rationalized by the fact that CoA alone, unlike LPS alone, is not a strong activator of NFκB as measured via NFκB luciferase assays (Fig. S4B). This supports a model where CoA alone primarily enhances a metabolic-epigenetic pathway, but not a SDTF activation pathway, that is sufficient to enhance basal *Il1b* and *Il1a* histone acetylation and transcription (Fig. 4A). Importantly, H3K27ac at the promoters of the expressed, non-TLR4 target genes *Nr1h2* and *Cox5a*, whose expression is not increased by CoA (Fig. 1D), was not increased by CoA treatment (Fig. S4C). This demonstrates that the acetylation status of specific (and not all) macrophage *cis*-regulatory elements are sensitive to CoA’s enhancement of mitochondrial glucose oxidation.

To further interrogate this specificity, we next used a more quantitative, unbiased Cut-and-Tag (CnT) approach to assess genome-wide H3K27ac changes in response to CoA alone, LPS alone, or CoA/LPS co-treatment. H3K27ac CnT faithfully captured macrophage regulatory regions identified in previous ChIP-seq studies with improved signal-to-noise and less background (Fig. S4D). Of 116,083 H3K27ac CnT peaks identified across all experimental samples, only 4869 (4.19% of total) peaks were significantly increased in H3K27ac CnT read density by CoA treatment alone versus no treatment, further supporting the notion that exogenous CoA influences histone acetylation not globally, but at select *cis*-regulatory regions (Fig. 4D). Likewise, LPS treatment increased H3K27ac at just 9034 peaks (7.78% of total), demonstrating the LPS response is restricted to specific *cis*-regulatory regions as well (Fig.4D). The H3K27ac peaks increased by CoA alone or LPS alone showed highly significant overlap that wouldn’t occur by chance, as nearly 50% (2432) of CoA-increased H3K27ac peaks were shared with LPS-increased H3K27ac peaks (Fig.4E). Gene Ontology (GO) analysis of genes associated with these 2432 CoA- and LPS-increased H3K27ac peaks revealed a strong enrichment for categories such as inflammation, cytokine production, and leukocyte activation (Fig. 4F), as these peaks either were in/adjacent to loci whose transcription we already had validated is positively influenced by CoA (e.g. *Il1b*), or in putative CoA-regulated proinflammatory genes (e.g. *Ccl5*) (Fig. S4E). HOMER *de novo* motif enrichment analysis revealed the 2432 CoA- and LPS-increased H3K27ac peaks were significantly enriched for NFκB and PU.1 binding sites (Fig. 4G), supporting our model (Fig. 2A) that both CoA and LPS positively influence histone acetylation primarily at sites of signal-regulated TF binding to developmentally-established macrophage enhancers. Looking quantitatively at global trends in behavior of these 2433 peaks, their low existing H3K27ac levels in untreated macrophages were strongly increased by CoA alone, more strongly increased by LPS, and showed an even further increased by CoA/LPS co-treatment (Fig. 4H). This H3K27ac responsiveness hierarchy (CoA/LPS > LPS > CoA > no treatment) exactly matches the transcriptional responsiveness hierarchy for genes such as *Il1b* and *Il1a* to the same treatments (Fig. 1B), providing a strong correlation between the strength of influence on histone acetylation by each stimulus, and the strength of proinflammatory gene induction. This H3K27ac responsiveness hierarchy was present at *Il1b* and *Il1a* regulatory regions, and revealed to exist at putative regulatory regions for other proinflammatory genes, such as *Il12b, Ccl5*, and *Cd40*. (Fig. 4I, Fig. S4E). In contrast to these effects on H3K27ac, NFκB luciferase experiments did not reveal enhanced NFκB activation upon CoA/LPS co-treatment versus LPS alone (Fig. S4B). Thus, our data supports our model (Fig. 2A) where CoA/LPS co-treatment produces a higher-in-magnitude transcriptional response than LPS alone via CoA-enhanced metabolic-epigenetic support of histone acetylation that enables and drives maximal transcription of specific proinflammatory genes.

### Tonic TLR signaling regulates basal macrophage glucose uptake, metabolic homeostasis, histone acetylation, and gene expression

Coenzyme A is not itself a nutrient or carbon source, but rather a metabolic co-factor that supports the movement of carbon from various sources (e.g. glucose) into metabolic pathways (e.g. the TCA cycle) and pathway endpoints (e.g. histone acetylation that positively influences gene expression). While increasing intracellular CoA alone might expand the capacity of a macrophage to carry out these processes and tune-up proinflammatory gene expression via increasing basal histone acetylation, such an effect would still be dependent on a cellular growth factor signal that drives glucose acquisition, metabolism, and utilization for histone acetylation. In *in vitro* models, strong TLR agonism with exogenous PAMPs like LPS is an anabolic signal that instantaneously increases macrophage glucose uptake (*21*, *54*) and oxidation for histone acetylation (*22*, *23*). However, BMDM cultures contain both PAMPs (e.g. LPS from serum), and dying cells that modulate the outputs of bystander macrophages exposed to exogenous stimuli (e.g. exogenous IL-4 added to BMDMs can only activate certain tissue repair gene expression programs if apoptotic cells are present as a co-signal (*55*)). Thus, both PAMPs and DAMPs released from stressed/dying cells represent a “tonic” TLR signal source in *in vitro* models. Indeed, others have reported that even in the absence of exogenously added PAMPs like LPS, both *Myd88*^−/−^ and *Ticam1*^−/−^ single knockout BMDMs show impaired basal glucose uptake (*23*). Inspired by this observation, we set out to provide evidence for the existence of tonic TLR signaling, and demonstrate its importance of TLRs in regulating the basal nutrient uptake, metabolic, epigenetic, and gene expression state of macrophages (Fig. 5A).

**Figure 5.**
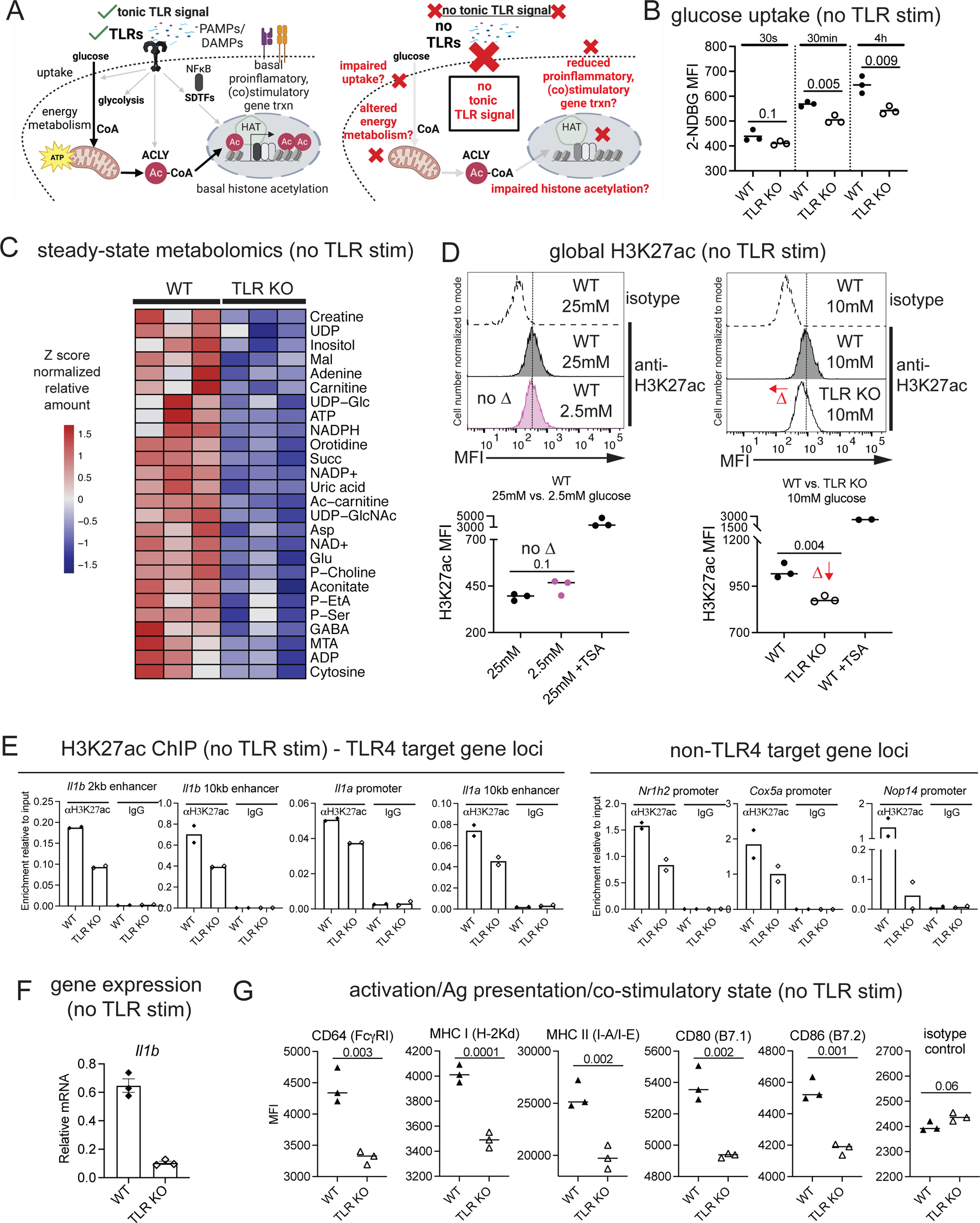
Tonic TLR signaling regulates basal macrophage glucose uptake, metabolic homeostasis, histone acetylation, and gene expression. **A**. Model depicting predicted regulation of macrophage biology by tonic TLR signaling. **B.** 2-NDBG uptake in WT vs. TLR KO rBMDMs. n=3, mean is shown, numbers above bars are Student’s t-test P values (planned comparisons). **C.** For approximately 130 metabolites detected, Z score-normalized steady-state levels of 26 metabolites displaying significant differences (all decreased, p<0.05, One-Way ANOVA) between WT and TLR KO rBMDMs. n=3 biological replicates per genotype. **D.** Left – H3K27ac ICS-flow cytometry in rBMDMs cultured in DMEM with 25mM versus 2.5mM glucose. Top histograms representative of data quantified below. Right - H3K27ac ICS-flow cytometry in WT vs. TLR KO supBMDMs cultured in RPMI with 10mM glucose. Top histograms representative of data quantified below. n=3, mean is shown, and numbers above bars are P values from Student’s t-test (planned comparisons). TSA-treated cells represent H3K27ac positive control. **E.** H3K27ac levels at *cis*-regulatory regions in *Il1b* and *Il1a* loci (left) and non-TLR4 target gene promoters (right) assessed by ChIP-qPCR in WT versus TLR KO supBMDMs. Mean is shown, each data point represents an independent ChIP experiment/biological replicate (n=2 per condition). IgG control ChIP-qPCR shows specificity of H3K27ac signal. **F.** Basal *Il1b* expression in WT versus TLR KO rBMDMs. n=3, mean ± s.e.m. is shown. **G.** Cell-surface expression of the indicated markers measured by flow cytometry on WT versus TLR KO supBMDMs. n=3, mean is shown, numbers above bars are Student’s t-test P values (planned comparisons).

While Myd88^−/−^ Ticam1^−/−^ double KO BMDMs are generally thought to be deficient in all TLR signaling, other TLR-proximal signaling molecules that regulate macrophage nutrient uptake and metabolism have been described, such as BCAP (*7*). Thus, we chose to interrogate tonic TLR signaling using BMDMs from *Tlr2*^−/−^ *Tlr4*^−/−^ *Unc93b1*^−/−^ mice, which lack functional signaling from all cell surface and endosomal TLRs (henceforth referred to as TLR KO BMDMs)(*56*). We found that similar to single KO Myd88^−/−^ and Ticam1^−/−^ BMDMs, TLR KO rBMDMs and supBMDMs showed impaired basal glucose uptake compared to wildtype (WT) BMDMs in the absence of exogenously added TLR ligands (Fig. 5B, Fig. S5A). This supports the existence of tonic TLR signals in the form of PAMPs/DAMPs in BMDM cultures that drive basal glucose uptake. To assess the consequences of this impaired glucose uptake on macrophage metabolism, we performed polar metabolomics on WT and TLR KO rBMDMs. This revealed a number of significant (P<0.05) decreases in the steady-state levels of metabolites important for energy metabolism and redox homeostasis in rBMDMs lacking TLRs, including ATP, NAD, NADPH (Fig. 5C). Notably, acetyl-CoA levels were decreased in TLR KO macrophages by approximately 45% (P=0.06, Fig. S5B). Thus, the impaired glucose uptake in TLR KO macrophages due to lack of tonic TLR signals is linked to significant changes in the cellular metabolic state.

We next sought to connect these defects in glucose acquisition and metabolism in the absence of tonic TLR signaling to changes in histone acetylation. In other cell culture systems where anabolic signals are present, such cancer cells exposed to serum growth factors, and adipocytes exposed to insulin, extracellular glucose concentrations dictate glucose-derived acetyl-CoA production and histone acetylation in an ACYL-dependent fashion (*57*). Interestingly, we did not detect any differences in global H3K27ac in rBMDMs cultured in low (2.5mM) versus high (25mM) glucose, suggesting that unlike other cell types, altering environmental glucose concentrations is not sufficient to not influence basal histone acetylation in the presence of an anabolic tonic TLR signal (Fig. 5D, left). However, upon quantifying global H3K27ac in WT versus TLR KO supBMDMs both cultured in 10mM glucose, we detected significantly reduced global H3K27ac in the absence of TLRs (Fig. 5D, right). This demonstrates that tonic TLR signaling is a critical regulator of not only basal glucose uptake and energy metabolism, but also of the basal histone acetylation state of the macrophage epigenome. We confirmed decreases in histone acetylation in the absence of tonic TLR signaling by H3K27ac ChIP in supBMDMs at regulatory regions in the *Il1b* and *Il1a* loci (Fig. 5E, left). Surprisingly, these decreases in basal histone acetylation extended to non-TLR4 target gene promoters (Fig. 5E, right), supporting the flow cytometry result that the absence of tonic TLR signaling has far-reaching, global effects on the H3K27ac macrophage epigenome that goes beyond TLR-regulated host defense genes.

Finally, we sought to connect the observed changes in basal histone acetylation in the absence of tonic TLR signaling to changes in basal macrophage gene expression and activation state. Consistent with reduced H3K27ac at *Il1b* regulatory regions, qPCR revealed reduced basal *Il1b* expression in TLR KO versus WT rBMDMs (Fig. 5F). TLRs are well-known to regulate the expression of genes/proteins critical for macrophage phagocytic and cytotoxic function (e.g. Fc receptors), as well as expression of antigen presentation (e.g. MHC class I and II), and co-stimulatory (e.g. B7.1, B7.2) molecules important for activation of T cell-mediated adaptive immunity. Indeed, cell surface levels of these molecules was significantly reduced on TLR KO versus WT rBMDMs (Fig. 5G), demonstrating that tonic TLR signaling maintains their basal expression levels. Taken together, these data represent clear evidence of changes in basal glucose uptake, metabolism, epigenetics, and gene/protein expression, in TLR KO macrophages in the absence of exogenous TLR agonists. Thus, tonic TLR signaling is an anabolic nutrient acquisition and growth factor signal that shapes multiple interconnected metabolic-epigenetic aspects of the macrophage phenotype (Fig. 5A).

### Exogenous CoA’s effects on histone acetylation and proinflammatory gene transcription are dependent on tonic TLR signaling

Having demonstrated the existence of tonic TLR signaling that acts as a nutrient/glucose acquisition and utilization signal supporting macrophage histone acetylation, we next sought to test the hypothesis that exogenous CoA’s effects on proinflammatory gene expression and histone acetylation depend on this tonic TLR signal (Fig. 2A). This hypothesis stemmed from the observation that CoA and strong TLR signaling induced by the exogenous PAMP LPS demonstrated clear synergistic effects on proinflammatory gene induction relative to either treatment alone in both naïve (Fig. 1B) and tolerized (Fig.1C) BMDMs. Moreover, we also observed while neither LPS or CoA alone could individually overcome the immunosuppressive effects of 4-OHE1 on *Il1b* induction, combined LPS/CoA synergized to rescue *Il1b* induction (Fig. 6A). Thus, based on multiple observations of synergy, we hypothesized that the ability of CoA alone to enhance proinflammatory gene transcription would be dependent on tonic TLR signaling. Indeed, we found the ability of CoA treatment alone to transcriptionally upregulate *Il1b*, *Il1a*, and *Nos2* expression was almost completely lost in TLR KO BMDMs (e.g. <1% of the *Il1b* fold induction observed in WT BMDMs)(Fig. 6B, Fig. S6A). Thus, CoA’s ability to upregulate proinflammatory gene transcription is largely TLR-dependent.

**Figure 6.**
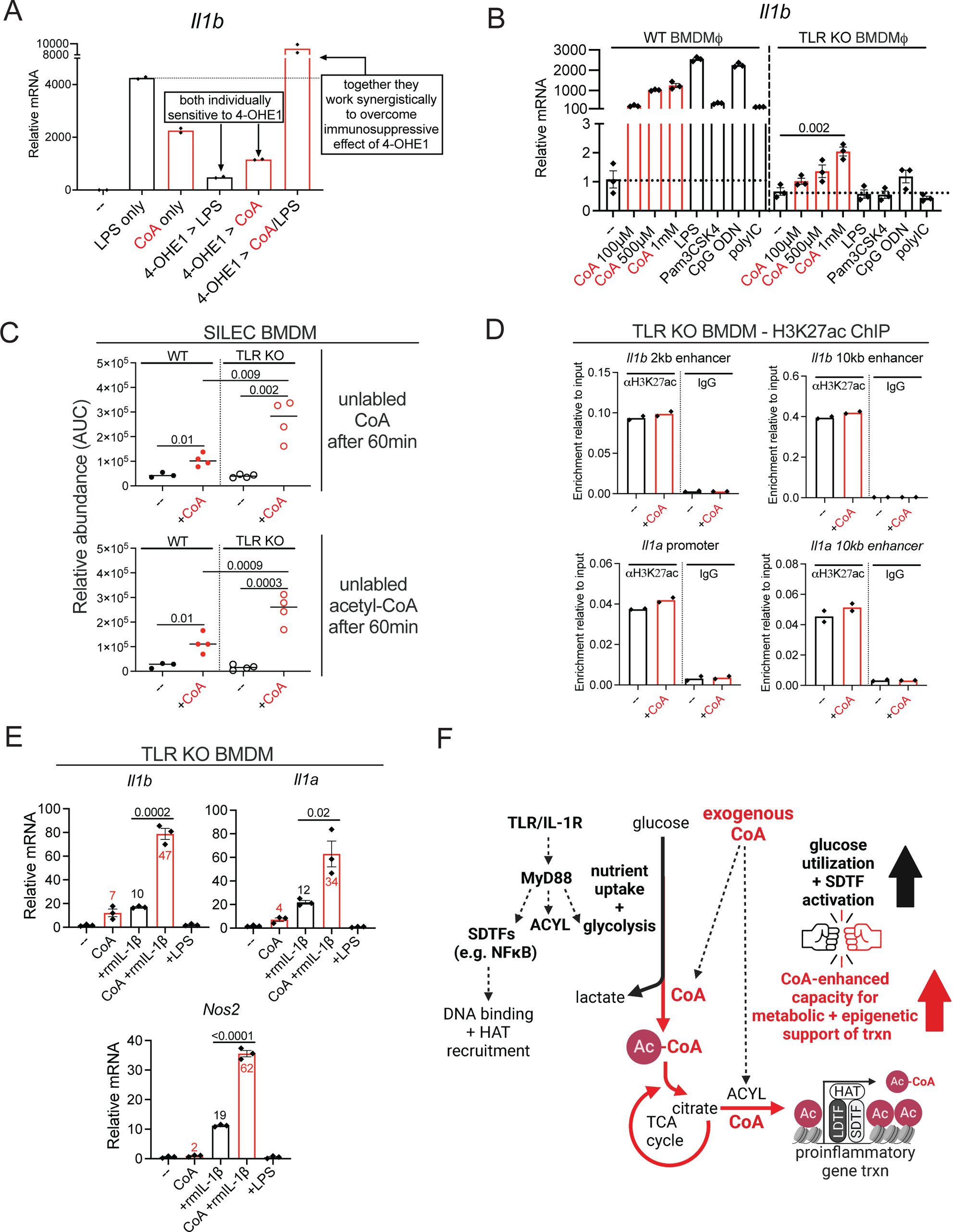
Exogenous CoA’s effects on histone acetylation and proinflammatory gene transcription are dependent on tonic TLR signaling. **A**. Il1b qPCR in rBMDMs treated with LPS alone, CoA alone (250μM), or CoA/LPS co-treatment, without or with 4-OHE1 (5μM) pretreatment. n=2, mean is shown. **B.** *Il1b* qPCR in WT and TLR KO BMDMs treated with CoA or TLR ligands for 4h. n=3, mean ± s.e.m. is shown. **C.** Relative abundance of unlabeled CoA (top) and unlabeled acetyl-CoA (bottom) in WT versus TLR KO SILEC rBMDMs treated with unlabeled CoA (500μM) for 60 minutes. n=3, mean is shown, numbers above bars are Student’s t-test P values (planned comparisons). **D.** H3K27ac levels at *Il1b* and *Il1a cis*-regulatory regions as assessed by ChIP-qPCR in control or CoA-treated (500μM, 1.5h) TLR KO supBMDMs. Mean is shown, each data point represents an independent ChIP experiment/biological replicate (n=2 per condition). IgG control ChIP-qPCR shows specificity of H3K27ac signal. **E.** *Il1b* qPCR in WT and TLR KO BMDMs pretreated with 500μM CoA for 2h before stimulation with 50ng/mL recombinant mouse IL-1β for 4h. **F.** Model depicting how exogenous CoA enhances macrophage mitochondrial glucose oxidation and acetyl-CoA production capacity, cooperating with TLR signaling to provide metabolic-epigenetic support of proinflammatory gene transcription.

We initially hypothesized that the inability of CoA to upregulated proinflammatory gene transcription in TLR KO BMDMs might reflect the inability of these cells to take up extracellular CoA, as TLR signaling drives macropinocytosis (*58*) that can mediate acquisition of extracellular molecules supporting anabolic metabolism (*59*). However, like WT SILEC rBMDMs, TLR KO SILEC rBMDMs were able to take up unlabeled exogenous CoA (Fig. 6C, top), ruling out any impairment in CoA uptake in the absence of tonic TLR signaling (which we found had no effect on macropinocytotic FITC-dextran uptake, Fig. S6B)). We next hypothesized that since TLR KO rBMDMs have impaired glucose uptake and altered metabolism, that perhaps the utilization of this newly-acquired CoA for glucose-derived acetyl-CoA production might be impaired in TLR KO rBMDMs. However, TLR KO SILEC rBMDMs actually had increased steady-state levels of unlabeled acetyl-CoA versus WT SILEC BMDMs (Fig. 6C, bottom). We cannot be certain, however, whether the carbon sources utilized to produce this acetyl-CoA (e.g. glucose, acetate, fatty acids, etc.) differ between WT and TLR KO BMDMs since we are only tracing unlabeled CoA (and not acetyl group carbons) in this experimental set-up.

A steady-state increase in a metabolite could be because 1) its production is increased with no change in consumption, 2) its production is unchanged but its consumption is reduced, or 3) some combination of these changes. Thus, we hypothesized that the increase in steady-state unlabeled acetyl-CoA levels in TLR KO versus WT SILEC rBMDMs could be because the unlabeled acetyl-CoA being produced is not being utilized for a downstream process required for increasing proinflammatory gene expression – namely, histone acetylation at *cis*-regulatory elements in proinflammatory gene loci. Thus, we tested whether CoA supplementation, which was sufficient to increase histone acetylation in WT BMDMs (Fig. 4C), could do the same in TLR KO BMDMs. Indeed, H3K27ac ChIP revealed that histone acetylation at *cis*-regulatory elements in the *Il1b* and *Il1a* loci was not increased by CoA supplementation in TLR KO BMDMs (Fig. 6D), consistent with an inability of CoA to upregulate *Il1b* and *Il1a* in these cells. This supports a model whereby TLR KO BMDMs can take up extracellular CoA and use it for acetyl-CoA biosynthesis, but this acetyl-CoA is not able to be utilized for histone acetylation that increases basal proinflammatory gene expression.

Finally, we reasoned that if CoA’s ability to upregulate proinflammatory gene expression was truly dependent on synergy and cooperation with tonic TLR signaling, then CoA should be able synergize with other TLR-like signals to enhance proinflammatory gene transcription even in the absence of TLRs. Like TLRs, the Interleukin 1 receptor (IL-1R) signals via MyD88 to upregulate proinflammatory gene expression in response to IL-1 cytokines. MyD88 signaling both activates SDTFs, and MyD88-TBK1-driven glycolysis (*54*) that provides a source of glucose-derived acetyl-CoA for histone acetylation. Indeed, we found that CoA synergized with recombinant IL-1β to significantly upregulate proinflammatory gene transcription in TLR KO BMDMs where no tonic TLR signal is present (Fig. 6E). This result definitively rules out the possibility that the proinflammatory effect of CoA are solely due to contaminating endotoxin, or other TLR ligands, in our yeast-derived CoA preparations. It also refutes the hypothesis that CoA is proinflammatory because it is a direct TLR ligand as has been suggested (*60*), as no TLRs are present in this experimental set-up in which CoA and recombinant IL-1β/MyD88 activation synergize to produce a proinflammatory effect. This result further supports a model in which the proinflammatory effects of CoA on WT macrophages lie in its ability to synergize with TLR signaling, acting as a direct metabolic adjuvant that provides enhanced capacity for metabolic-epigenetic support of histone acetylation and proinflammatory gene transcription (Fig. 6F).

### CoA enhances TLR agonist anti-tumor activity

As *in vivo* murine proinflammatory responses to LPS are largely macrophage-dependent (*61*, *62*), we tested if CoA could act as a metabolic adjuvant *in vivo* and enhance this response. CoA alone showed minimal proinflammatory activity in naïve mice as assessed by serum cytokine levels following CoA injection (Fig. 7A). However, levels of nearly two-thirds of the LPS-induced proinflammatory cytokines, chemokines, and growth factors quantified were significantly enhanced by CoA/LPS co-administration versus LPS alone (Fig. 7A). Thus, the ability of CoA to act as a metabolic adjuvant and enhance LPS- and TLR4-driven proinflammatory responses *in vitro* extends to mice *in vivo*.

**Figure 7.**
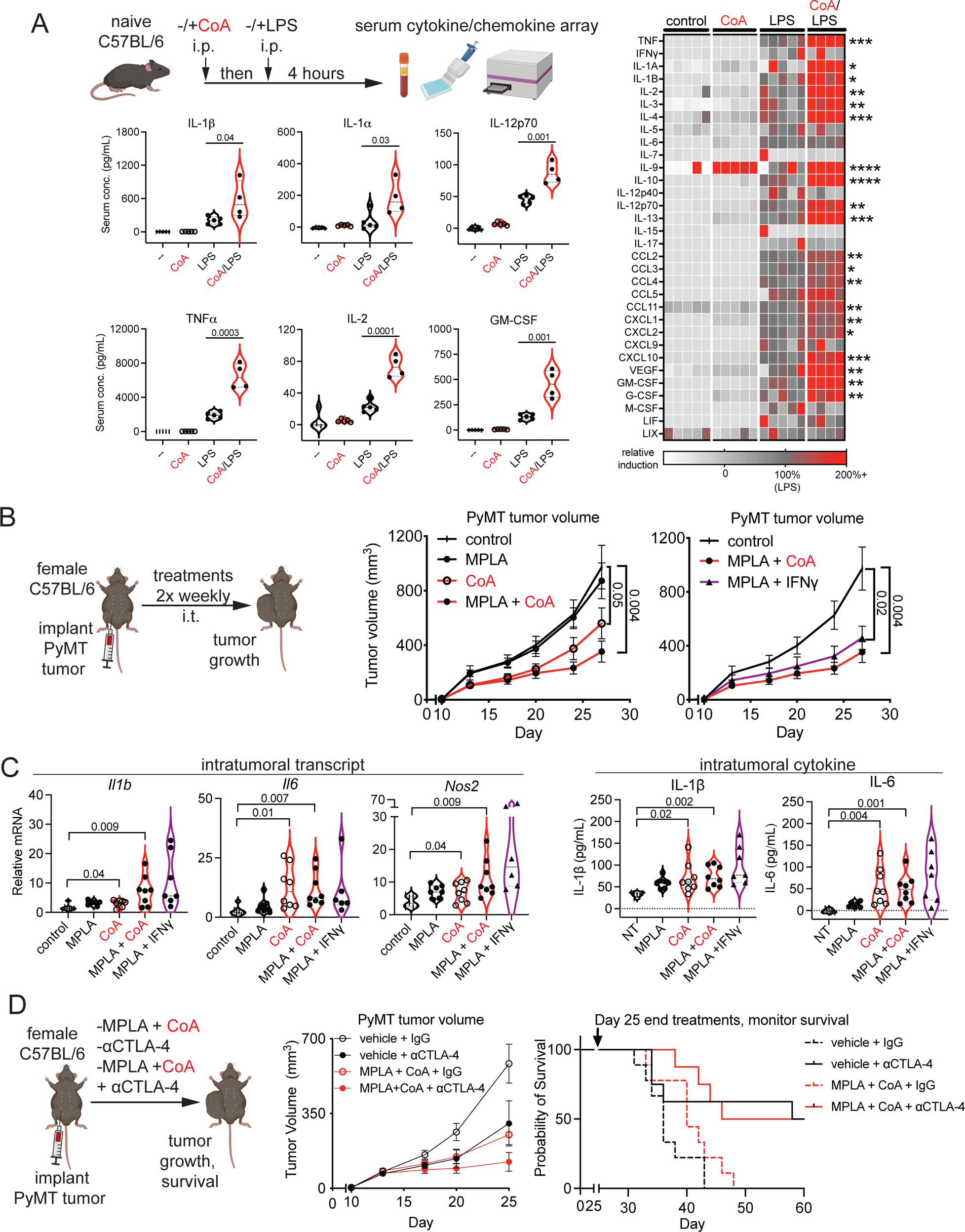
CoA enhances TLR agonist anti-tumor activity. **A**. Serum levels of select proteins in male C57BL/6 mice intraperitoneally (i.p.) injected with CoA (250 mg/kg), LPS (3 mg/kg), or CoA/LPS. Violin plots for select serum proteins (left), and heatmap showing relative induction of all cytokines, chemokines, and growth factors quantified (right), with induction in “LPS” group set to 100%. For violin plots, numbers above bars are P values from Student’s t-test for the indicated conditions (planned comparisons). For heatmap, *P<0.05. *P<0.01, ***P<0.001, and ****P<0.0001 between “LPS” and “CoA/LPS” groups in each row (Student’s t-test, planned comparisons). “x” denotes protein for which at least one sample in the “CoA/LPS” group exceeded the linear range of the standard curve, underestimating induction in this group. n=5 mice per group except “CoA/LPS”, n=4. **B.** Orthotopic MMTV-PyMT breast tumor volume in female C57BL/6 mice. Left – volume in control tumors versus tumors treated intratumorally (i.t.) with MPLA, CoA, or MPLA+CoA. Right – volume in control, MPLA+CoA-treated, and MPLA+IFNγ-treated tumors. Numbers represent Student’s t-test P values at Day 27 (planned comparisons). n=8 mice per group. **C.** Intratumoral gene expression (left) and cytokine levels (right) from mice in **A**. 16 hours after final i.t. treatment. n=7 or 8 mice per group. Numbers represent Student’s t-test P values (planned comparisons). **D.** Orthotopic MMTV-PyMT breast tumor volume (left) and survival (right) in female C57BL/6 mice treated for 25 days with controls, CoA, αCTLA-4, or CoA/αCTLA-4 combination therapy. n=10 mice per group.

We tested whether CoA supplementation could improve outcomes in situations where macrophage tolerance contributes to impaired host defense. The use of immunostimulatory agents such as TLR agonists to reprogram tumor-associated macrophages (TAMs) to a proinflammatory phenotype that supports anti-tumor immunity has been extensively studied (*63*), but with limited clinical success (*64*). The ineffectiveness of TLR agonists as monotherapies may lie in tolerance. Similarities have been noted between tolerized macrophages and immunosuppressive TAMs (*65–67*), suggesting DAMPs released from stressed/dying cells in the tumor microenvironment (TME) chronically stimulate TLRs, conditioning TAMs towards tolerance (*68*). Breast cancer TAMs are a potent source of TGFβ (*69*), a hallmark cytokine of tolerized macrophages (*5*). Moreover, TAMs isolated from spontaneous MMTV-PyMT breast tumors display tolerance, as the TLR4 agonist monophosphoryl Lipid A (MPLA, a LPS derivative and clinically-approved vaccine adjuvant) is unable to upregulate proinflammatory genes involved in anti-tumor immunity *in vitro,* or slow tumor growth *in vivo*, unless combined with IFNγ (*12*). Interestingly, CoA’s effects described thus far in many ways phenocopy IFNγ (a.k.a. “macrophage-activating factor”), a cytokine known to potentiate LPS-dependent macrophage proinflammatory responses (often called “priming”) (*70*, *71*), reverse macrophage tolerance (*72–75*), and promote macrophage anti-tumor activity (*76*, *77*). These findings led us to hypothesize that CoA co-administration with MPLA would unlock the anti-tumor activity of this TLR agonist.

Similar to other tumor models (*12*, *78*, *79*), intratumoral (i.t) MPLA alone had no effect on MMTV-PyMT orthotopic breast tumor growth (Fig. 7B left, Fig. S7A). However, an MPLA + CoA combination therapy significantly slowed tumor growth (Fig. 7B left, Fig. S7A), phenocopying MPLA+IFNγ (Fig. 7B right, Fig. S7A). Interestingly, CoA alone had a measurable effect on tumor growth (Fig. 7B left, Fig. S7A), suggesting CoA might help TAMs overcome tolerance and enable proinflammatory responses to endogenous DAMPs. In support of this, both CoA monotherapy, and MPLA + CoA, were superior to MPLA monotherapy at elevating i.t. proinflammatory gene expression and cytokine levels (Fig. 7C). These measurements were made 16 hours after the final i.t. therapy injection, a timepoint when an acute inflammatory response would have resolved, suggesting CoA enables sustained TAM proinflammatory responses to endogenous DAMPs and/or MPLA. Interestingly, both CoA and IFNγ prevented an MPLA-induced increase in intratumoral lactate levels (Fig.S7B). Moreover, the anti-tumor effects of the MPLA + CoA combination therapy was dependent on TAMs, as their depletion abrogated its ability to slow tumor growth (Fig. S7C-D).

Finally, when combined with T cell-targeted checkpoint blockade therapy, MPLA + CoA had additive affects with anti-CTLA4 on reducing tumor aggression as measured by PyMT tumor growth suppression during the first 25 days post-tumor implantation (Fig. 7D, left). Upon discontinuation of treatments at day 25, both MPLA + CoA alone, and MPLA + CoA in combination with anti-CTLA-4, initially showed promise in extending median survival compared to appropriate controls between days 35 and 35 (Fig. 7D, right). However, this effect waned with time, and MPLA + CoA did not increase the number of mice receiving checkpoint blockade that rejected tumors (Fig. 7D, right). This suggests that unlike checkpoint blockade, continued MPLA + CoA treatments might be required to produce sustained anti-tumor growth effects and improve survival in this tumor model.

Collectively, these results support the proposed hypothesis, and suggest that CoA-dependent tolerance reversal improves efficacy TLR agonist immune therapy via enabling TAM proinflammatory reprogramming. Moreover, they demonstrated the potential of a CoA + TLR agonist combination therapy to work together with checkpoint blockade immunotherapy to reduce tumor aggression.

### CoA enhances macrophage anti-microbial activity against *Legionella pneumophila* via CoA-supported itaconate biosynthesis

Evidence suggests that tolerance contributes to host immunosuppression during infection, particularly with pathogens that replicate within macrophages. Intracellular bacterial pathogens generally suppress macrophage mitoOXPHOS, shifting host metabolism towards aerobic glycolysis (*80*), a shift identical to that observed metabolically during LPS tolerance. This suggests TLR signaling triggered by chronic PAMP exposure during infection, pathogen effectors that disrupt mitoOXPHOS (*81*), or both, promote a mitoOXPHOS/aerobic glycolysis shift to create a tolerized replicative niche for pathogens. IFNγ enhances macrophage anti-microbial activity (*82*, *83*) via induction of host genes that restrict intracellular pathogen growth, with full induction of restriction genes such as *Nos2* requiring synergy between IFNγ and TLR signaling (*76*). Thus, IFNγ’s restrictive effects may lie in its ability to reverse tolerance and enable full expression of host restriction genes. Given CoA’s ability to (re)direct glucose into the mitochondrial TCA cycle and reverse tolerance, we hypothesized that CoA would phenocopy IFNγ in restricting intracellular pathogen replication in macrophages.

To test this, we utilized *Legionella pneumophila*, a gram-negative intracellular bacterial pathogen and causative agent of a severe type of pneumonia, Legionnaires’ Disease. During macrophage infection, *L. pneumophila* triggers multiple TLRs (*84*), and secretes multiple effectors that cause mitochondrial stress and suppress mitoOXPHOS (*85–87*). This diverts host metabolism towards aerobic glycolysis (*85*), indicative of tolerance. IFNγ potently restricts *L. pneumophila* intracellular replication in a manner that depends on host induction of six antimicrobial defense genes, including *Nos2* and *Irg1* (*88*). Basal expression of these 6 genes was increased in uninfected BMDMs by CoA alone, and induction was further enhanced with CoA/LPS cotreatment for some genes (Fig. S8A), prompting us to test the whether CoA could restrict *L. pneumophila*. To track *L. pneumophila* replication in host macrophages, we used a previously validated luminescent strain (*89*, *90*). As *L. pneumophila* flagellin potently triggers macrophage pyroptosis, we used a flagellin-deficient (Δ*flaA*) luminescent strain to avoid this confounding effect (*89*, *90*). Real-time tracking demonstrated robust Δ*flaA L. pneumophila* replication in BMDMs (Fig. 8A). However, CoA supplementation restricted growth in a dose-dependent manner (Fig. 8A, left), phenocopying IFNγ (Fig. 8A, right). CoA-mediated restriction, which was confirmed at the level of bacteria colony-forming units (CFUs, Fig. S8B), occurred independent of direct effects on *L. pneumophila* growth or luminescence (Fig. S8C), or BMDM viability (Fig. S8D).

**Figure 8.**
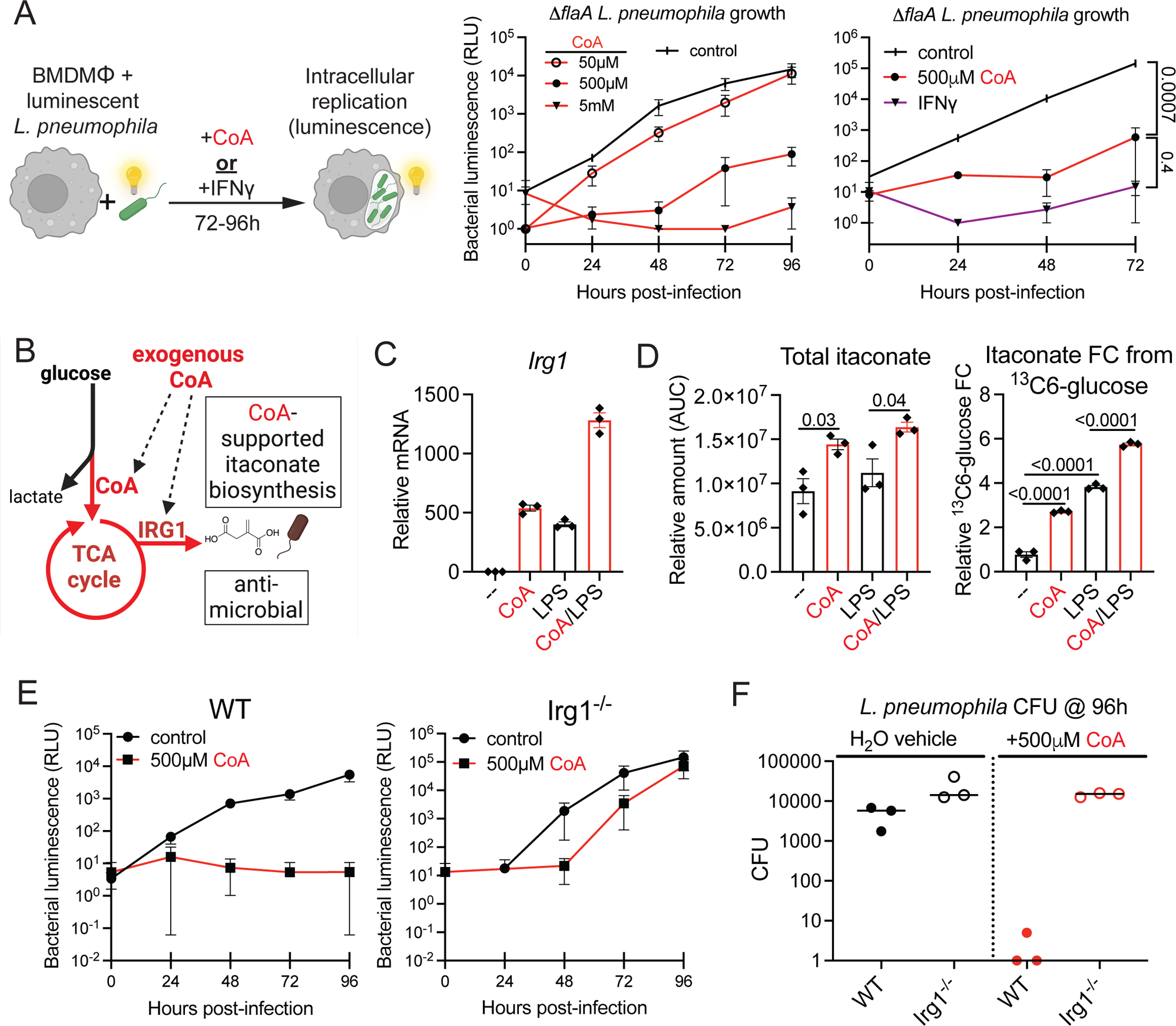
CoA enhances macrophage anti-microbial activity against *Legionella pneumophila* via CoA-supported itaconate biosynthesis. **A**. Luminescent Δ*flaA Legionella pneumophila* intracellular growth in infected supBMDMs. Left – intracellular bacterial replication over time in control or CoA-treated supBMDMs. Right - intracellular bacterial replication over time in supBMDMs treated with CoA or IFNγ (6ng/mL). Numbers represent Student’s t-test P values (planned comparisons). n=3 biological replicates per condition. **B.** Model depicting how CoA could support itaconate production via 1) increasing metabolic-epigenetic support of *Irg1* expression, and 2) supporting increased flux of glucose carbon into the TCA cycle. **C.** *Irg1* qPCR in supBMDMs treated with CoA (250μM), LPS, or CoA/LPS for 2h. **D.** Total steady-state itaconate levels (left), and relative ^13^C_6_-glucose fractional contribution (FC) to itaconate (right), in rBMDMs treated with CoA (250μM), LPS, or CoA/LPS for 2h. **E.** Luminescent Δ*flaA Legionella pneumophila* intracellular growth over time in control or CoA-treated supBMDMs from WT or *Irg*1^−/−^ macrophages. **F.** *L. pneumophila* colony forming units (CFUs) from infected WT and *Irg*1^−/−^ supBMDMs in **E** at 96h.

Itaconate is a host TCA cycle-derived anti-microbial metabolite induced by inflammatory cues that is directly secreted from mitochondria into pathogen-containing compartments to restrict pathogen growth via its reactive electrophilic properties. Increased itaconate biosynthesis from glucose-derived carbon during infection depends on both the transcriptional induction of the *Irg1* gene by immunostimulatory cues (e.g. TLR signaling), and enhanced flux of glucose-derived carbon into the mitochondrial TCA cycle from which itaconate is synthesized from aconitate. As CoA could simultaneously positively influence both of these processes (Fig. 8B), we further investigated the role of itaconate and *Irg1* in *L. pneumophila* restriction by CoA. As mentioned, CoA was sufficient to induce *Irg1*, and also enhanced LPS-induced *Irg1*, fulfilling the criteria regarding gene induction (Fig. 8C, Fig. S8A). Examining our metabolomics data, CoA was sufficient to increase both total itaconate levels, and itaconate biosynthesis from glucose-derived carbon (Fig. 8D), an effect that both was enhanced upon CoA/LPS co-treatment. This effect phenocopies LPS/IFNγ co-treatment (Fig. S8E). Based on these data, we tested whether CoA’s restriction of *L. pneumophila* might be *Irg1*-dependent. Indeed, compared to WT BMDMs, CoA’s restriction of *L. pneumophila* growth was impaired in *Irg1* KO BMDMs, with the data suggesting CoA’s effects are at least partially (if not largely) dependent on *Irg1* (Fig. 8E). Thus, this supports a model where CoA restricts *L. pneumophila* largely through coordinate transcriptional (i.e. acetyl-CoA dependent histone acetylation) and metabolic (i.e. mitochondrial glucose oxidation) support of *Irg1*-depednent itaconate biosynthesis. More broadly, these results support a view on chronic infection-induced tolerance whereby pathogen-driven mitoOXPHOS/aerobic glycolysis shifts promote decreased biosynthesis of proinflammatory and anti-microbial metabolites (acetyl-CoA, itaconate) to create hospitable replicative pathogen niche in tolerized macrophages (Fig. S8F).

## Discussion

Our findings show that the capacity for macrophages to engage the primary metabolic-epigenetic pathway supporting TLR-dependent transcriptional programing, mitochondrial glucose oxidation for acetyl-CoA production, can be augmented by direct CoA supplementation. Moreover, they demonstrate that macrophage tolerance, an immunosuppressed state of innate immune memory driven by chronic TLR signaling and enforced by declines in both mitochondrial glucose oxidation (*22*, *34*) and histone acetylation at TLR target genes (*35*), is plastic and can be reversed by direct CoA supplementation. The effects of CoA are all critically dependent on TLR signaling itself, as we show for the first time that in the absence of any tonic TLR “growth factor” signal, macrophage nutrient uptake, metabolism, epigenetic state, and gene expression are all significantly altered. Taken together with our previous study demonstrating the potent anti-inflammatory effects of CoA depletion (*34*), our work shows that effective levels of this key metabolic cofactor in macrophages governs their mitochondrial glucose oxidation capacity, and in turn, metabolic-epigenetic support capacity for TLR-dependent macrophage proinflammatory responses.

How extracellular CoA is taken up by macrophages to positively influence CoA-dependent glucose utilization is unclear, as there is no described CoA transporter/uptake mechanism (*45*). That extracellular CoA is proinflammatory also raises the possibility that CoA released into the extracellular space during tissue damage represents a previously unappreciated mitochondrial DAMP (*1*) promoting sterile inflammation not via stimulating TLRs/other receptors, but via augmenting a specific macrophage metabolic-epigenetic axis supports proinflammatory gene expression. Our data suggests degradation by ENPPs works against CoA’s proinflammatory effects (Fig. S2C), and would antagonize CoA-driven inflammation following injury. This would be analogous to how other immunostimulatory molecules (ATP, cyclic dinucleotides) are degraded by ENPPs to resolve inflammation, but work against anti-tumor immunity (*91*).

Exogenous CoA supports IL-4-driven macrophage alternative activation (*92*), which is a macrophage phenotype also dependent on ACLY-dependent histone acetylation (*93*), but driven by different SDTFs (e.g. STAT6) activating different target genes. Going forward, whether other cell types besides macrophages can directly take up intact CoA (or other acly CoAs), and whether the uptake mechanisms are similar or divergent, are outstanding questions of interest. Similar to our results, CoA supplementation augmented mitochondrial glucose oxidation in T cells, though CoA was not isotopically traced into T cells in this study due to the lack of a commercial source of labeled CoA (*94*). Studies using radiolabeled ^3^H-acetly-CoA in which atoms in the acetate group can be traced suggest cell types such a breast cancer cells (*95*) and myeloid precursors (*96*) can acquire the labeled acetate from the extracellular space, though uptake of the CoA moiety could not be assessed with this tracing strategy. The SILEC system described here, in which the endogenous CoA pool is isotopically labeled to allow pulse-chase studies of the fate of intact unlabeled CoA, circumvents the need for labeled CoA in tracing studies, and is readily applicable to other cell culture systems. Moreover, the SILEC system allows for both interrogation of CoA uptake capacity, and investigation as to whether exogenously acquired CoA is utilized in the same manner as endogenously-produced CoA with regards to use in different acyl CoA biosynthesis pathways within the cell. Such studies will be important in understanding the ability of different cells types in complex microenvironments such as tumors to uptake CoA, and how competition for this metabolic cofactor might influence processes such as cell growth, tumor aggression, and anti-tumor immunity.

Immune suppression is a barrier to the treatment of inflammatory diseases, as both sterile and non-sterile PAMP/DAMP-triggered inflammation have evolved strong negative feedback mechanisms to both protect the host, and transition responses to an injury-reparative phase. In solid tumors including breast cancer, this response promotes tumor growth, with TAMs polarizing towards a wound-healing, TGFβ-producing, collagen-biosynthetic, immunosuppressive phenotype (*97*). The nutrient fluxes in TAMs are likely incompatible with supporting a proinflammatory phenotype, as the biosynthetic goals of these macrophages (collagen production for repair vs. proinflammatory mediator production for defense) are divergent. While TAMs are copious consumers of glucose in the TME (*98*), if the flux of this glucose is more directed towards lactate (immunosuppressive) or serine (collagen biosynthesis), this would be counterproductive to proinflammatory responses, and consistent with the aforementioned TAM tolerance phenotype (*12*). In turn, failures of therapies aimed at reprograming TAMs to a proinflammatory phenotype may lie in failure to address the metabolic programming enforcing tolerance. Our work combining MPLA with CoA represents proof-of-concept that combining an immunostimulatory cue with a “metabolic adjuvant” that helps macrophages re-engage the key metabolic flux supporting proinflammatory gene expression can improve TAM-targeted immunotherapy efficacy. Our results also suggest CoA supplementation may hold promise in reinvigorating host immunity in chronic infections associated with immunosuppression, sepsis-associated immune suppression (often accompanied by secondary infections, and shares parallels with cancer-associated immunosuppression (*11*)), and aging-associated immune decline.

## Acknowledgments

We thank the following people for their helpful advice and discussion regarding the project, and/or assistance with animals, reagents, experimentation, and data analysis: Dr. Jonathan Lakins and Dr. Roger Oria-Fernandez (Weaver Lab, UCSF); Dr. Gregory Barton and Dr. Kathleen Pestal (Barton Lab, UC Berkeley); Dr. Jordan Price and Kristen Witt (Vance Lab, UC Berkeley); Dr. Kelly Kersten (Krummel Lab, UCSF); Varun Bhadkamkar and Dr. Shaeri Mukherjee (USCF); Dr. Darren Gay (CoALA Biosciences); the support team in the UCLA Metabolomics Center for the LC-MS data analysis.

## Funding

This work was supported by:

American Diabetes Association Postdoctoral Fellowship #1-19-PDF-058 to G.A.T.

1F32CA236156-01A1 (NIH F32 Postdoctoral Award), 5T32CA108462-15 (NIH T32), and Sandler Program for Breakthrough Biomedical Research (postdoctoral independence award) to K.M.T.

The Netherlands Organization for Scientific Research VENI grant 09150161910024 to J.D-A

R35 CA242447-01A1 to V.M.W.

R01CA259111 and R01GM132261 to N.W.S.

T32GM142606 to D.S.K.

This work used:

the UCLA Metabolomics Center, supported by NIH Instrumentation Grant S10 OD016387

the Parnassus Flow Cytometry CoLab (PFCC, RRID:*SCR_018206*), supported in part by the DRC Center Grant NIH P30 DK063720

## Author contributions

V.M.W. supervised the project. G.A.T. conceptualized the project. G.A.T., K.M.T., J.t.H., D.S.K., J.F., J.D-A., R.E.V., N.W.S., and V.M.W. procured funding and resources. G.A.T., K.M.T., J.t.H., E.A.N., C.K., J.D-A., R.E.V., and N.W.S. designed experimental methodology. G.A.T., K.M.T., D.S.K., J.t.H., I.B., E.A.N., C.K., J.D-A., R.E.V. and N.W.S. performed experiments and analyzed data. G.A.T. curated data, wrote the manuscript, and edited the manuscript with input from all co-authors.

## Competing interests

Greg A. Timblin and Joshua N. Farahzad are co-founders and own equity Inapill Inc.

## Data and materials availability

All raw data is available upon request & will be made available upon publication of the peer reviewed manuscript.

## Materials and Methods

### Animals

All experiments were approved by the UCSF Institutional Animal Care & Use Committee (IACUC) to ensure ethical use of animals (Protocol AN194983-01), and performed under the supervision of the UCSF Laboratory Animal Resource Center (LARC). C57BL/6J mice (strain #000664) were purchased from Jackson Labs. Animals were housed under standard conditions (20-23°C, 50% humidity, 12h/12h light/dark cycle). Bones from TLR KO and Irg1 KO mice were generous gifts from Dr. Gregory Barton and Dr. Russell Vance, respectively (both UC Berkeley).

### Cell culture

For all experiments except *L. pneumophila* infection model, bone marrow-derived macrophages (BMDMs) were prepared from 6-12-week-old female C57BL/6J mice by plating bone marrow in DMEM (GeneClone 25-500) supplemented with 10% fetal bovine serum (FBS, HyClone SH30071.03m, non-heat inactivated), penicillin/streptomycin (Gibco 15140-122), and 20ng/mL recombinant murine M-CSF (Shenandoah 200-08 or BioLegend 576404) in non-TC treated Petri dishes, refreshing media every 3 days. Cells were harvested by lifting with PBS with 4mM EDTA with gentle scraping, and plated in TC-treated 24-well plates at 1 x 10^5^ cells per well on days 6-10. For experiments looking solely at naïve macrophage LPS responses, 100 ng/mL LPS was used. BMDM tolerance was induced by stimulating naïve BMDMs for 24 hours with 100 ng/mL LPS (Sigma L3024). Following a media change and 2 hour recovery period, tolerized and control naïve macrophages were (re)stimulated with 10 ng/mL LPS. For CoA pre-incubation experiments, FBS heat-inactivation was carried out by incubating HyClone SH30071.03m at 56°C for 30 minutes. Then,1 part CoA was mixed with 9 parts PBS, media, non-heat-inactivated FBS, or heat-inactivated FBS, in a 1.5mL tube and incubated at 37°C for 3h. Afterwards, the entirety of each tube was added to BMDMs.

For *L. pneumophila* infection experiments, BMDMs were prepared by plating bone marrow in RPMI (Gibco 21870-076) supplemented with 2mM glutamine (Gibco 25030081), 10% FBS (Gibco 16140-071, heat inactivated), and 5% supernatant from 3T3 cells expressing macrophage colony-stimulating factor (generated in-house in Vance Lab). RAW 264.7 macrophages were cultured in DMEM with 10% FBS and pen/strep. THP-1 cells were cultured in RPMI 1640 (GeneClone 25-506) supplemented with 10% FBS and pen/strep (both from ATCC via the UC Berkeley Cell Culture Facility). MMTV-PyMT breast cancer cell line was cultured in 50/50 mix of DMEM and F12 (Gibco 11765-054) with 10% FBS and pen/strep.

### Chemicals

Coenzyme A (CoA) used in the majority of experiments was from CoALA (CA02, lithium salt), though we confirmed CoA from two other sources (Sigma C4780, Cayman 16147) produced similar results *in vitro*. Sigma CoA was also used in the *in vivo* LPS challenge experiment, and in the metabolomics in Figure 2B. (Figure S2C, S2D metabolomics used CoALA CoA). Acetyl-CoA (A2056), Pantothenate/Vitamin B5 (21210), LPS (L3024) and poly IC (P0913) were from Sigma. 3’-dephosphoCoA (27390) and adenosine 3’,5’-diphosphate (24265) were from Cayman. 4-phosphopantetheine (CX11340) was from Chiralix. MPLA (tlrl-mpls), Pam3CSK4 (tlrl-pms) and ODN (tlr1-1826-1) were from InvivoGen. Recombinant murine IFNγ was from PeproTech (315-05) or BioLegend (575304). Recombinant murine IL-1β was from BioLegend (575102).

### Quantitative real-time PCR (qPCR) for gene expression

Cells were lysed in Trizol (Invitrogen) and total RNA isolated via chloroform extraction according to manufacturer’s instructions using GlycoBlue (AM9515) to assist RNA precipitation. cDNA was prepared from approximately 250ng of total RNA with random hexamer priming using M-MLV Reverse Transcriptase (BioChain Z5040002-100K). qPCR was performed on an Eppendorf Mastercycler Realplex2 epgradient S qPCR machine using PerfeCTaSYBR Green FastMix (Quantabio 95072-05K). Primers (IDT) were designed to amplify exon-spanning sequences in target mRNAs using Primer-BLAST. All data is normalized to *Hprt* expression using the delta-delta Ct method.

Primer sequences (5’-3’ F, 5’-3’ R):

Mouse

*Hprt -* TCAGTCAACGGGGGACATAAA, GGGGCTGTACTGCTTAACCAG

*Il1b -* CGTGGACCTTCCAGGATGAG, CGTCACACACCAGCAGGTTA

*Il1a -* AACGTCAAGCAACGGGAAGA, GCTGATCTGGGTTGGATGGT

*Il6 –* CGATAGTCAATTCCAGAAACCGC, TGGTTGTCACCAGCATCAGTC

*Tnf -* CCCTCACACTCAGATCATCTTCT, GCTACGACGTGGGCTACAG

*Il12b -* TCTCCTGGTTTGCCATCGTT, GGGAGTCCAGTCCACCTCTA

*Nos2 -* GTTCTCAGCCCAACAATACAAGA, GTGGACGGGTCGATGTCAC

*Irg1 -* AGCAACTACTCCTGCCATCC, CAAGCCGTGAATCATACCAGC

*Cybb -* TGCTAACATGGGGAACTGGG, AGAGGAAGACATTCAGCCCC

*Casp4 -* TAGGCTACGATGTGGTGGTG, TGCTGTCTGATGTCTGGTGT

*Irgm1 -* CAGCCCAAACCGTAGAGGAC, CGAGCTGAACTGCTCAGAGG

*Irgm3 -* CTGAGCCTGGATTGCAGCTTT, TGAACTCTGTGGGTCTGCTCTA

*Nr1h2 -* CAGCTGCAGTGCAACAAACG, GCATCTCGGGACTGAGGGTC

*Cox5a* – GATGCTCGCTGGGTGACATA, GCTCAGGAACCAGATCATAGCC

*Nop14* – GACAAACGCTTTGGGGAGTAT, TCATGATATCGCTGTTGCTCCA

*Sdc1* – GTCGAGGATGGAACTGCCAAT, TGTGTTCTCCCCAGATGTTTCA

*Iah1* – GGAAAACCCAGTGGCCGTTA, GGTTTGCGCTGTACTCATCC

*Gga1* – TGGACCTCTTGGGGAAAACC, CTTACTCTGCAGGTCACGGA

Human

*HPRT -* TGCTGAGGATTTGGAAAGGGT, GGGCTACAATGTGATGGCCT

*IL1B -* GCAGCCATGGCAGAAGTACC, TGTTTAGGGCCATCAGCTTCAA

*IL1A -* CGGTTGAGTTTAAGCCAATCCAT, AAGGTGCTGACCTAGGCTTG

*IL6 -* AGTCCTGATCCAGTTCCTGC, CAGGCTGGCATTTGTGGTTG

*TNF -* CTGCACTTTGGAGTGATCGG, TGTCACTCGGGGTTCGAGA

*NOS2 –* TCTCAAGGCACAGGTCTCTTC, CAGATGTTCTTCACTGTGGGG

### Monocyte isolation and stimulation

PBMC isolation was performed by differential density centrifugation over Ficoll-Paque (GE Healthcare). Percoll isolation of monocytes was performed as previously described (*99*). Briefly, 150-200 x 10^6^ PBMCs were layered on top of a hyper-osmotic Percoll solution and centrifuged for 15 min at 580g. The interphase layer was isolated and cells were washed with cold PBS. Cells were re-suspended in RPMI medium Dutch modified (Invitrogen) supplemented with 50 µg/mL gentamicin, 2 mM Glutamax, and 1 mM pyruvate, and counted. An extra purification step was added by adhering Percoll-isolated monocytes to polystyrene flat bottom plates (Corning) for 1 h at 37°C; a washing step with warm PBS was then performed to yield maximal purity. 10^5^ monocytes were added to flat-bottom 96-well plates (Greiner). Cells were stimulated with RPMI or Escherichia coli LPS (Sigma-Aldrich) at a concentration of 10 ng/mL for 4h-incubation at 37°C. In the specific conditions described, CoA was supplemented at two doses (low: 500µM CoA, high: 5mM) 2 hours prior to LPS stimulation. Total RNA was prepared using NucleoSpin RNA Plus Mini kit (Macherey-Nagel) according to manufacturer’s instructions for cDNA synthesis and qPCR as described above.

### *In vivo* LPS challenge and serum cytokine array

8 week-old male C57BL/6 mice (Envigo, 5 mice per group) were injected intraperitoneally (i.p.) with either PBS vehicle or 250 mg/kg CoA (Sigma C4780). 15 minutes later, mice were injected i.p. with either PBS vehicle or sublethal 3 mg/kg LPS dose (Sigma L2880). 4 hours later, mice were bled for serum cytokine analysis using MILLIPLEX MAP Mouse Cytokine/Chemokine Magnetic Bead Panel (Millipore *MCYTOMAG-70K*-PMX) according to manufacturer’s instructions. One sample was excluded from the CoA/LPS group data as it was identified as an outlier by the ROUT method across multiple quantified cytokines.

### PyMT breast cancer model

6 week old female C57BL/6 mice were anesthetized and 250K MMTV-PyMT tumor cells (100μL total volume, 1:4 mix of cells in PBS:matrigel) injected into the inguinal fat pad near the 4^th^ mammary gland using a 1mL syringe and 26G needle (Day 0). Palpable tumors were detectable in some animals by Day 7, and all animals by Day 10. Twice-weekly intratumoral injections were initiated on Day 11 using an insulin syringe (BD Biosciences) to deliver 50uL volumes of MPLA (1mg/tumor, DMSO vehicle), CoA (5, 50, or 250 mg/kg, H_2_O vehicle), MPLA + CoA, or vehicle controls. Tumor volumes were measured with calipers starting on Day 13 prior to injections. At experimental endpoint, mice were sacrificed and tumors excised, weighed, and either immediately processed for flow cytometry, or flash frozen, homogenized, and stored at −80°C for other uses.

For TAM depletion experiment, 1mg of anti-CSF-1R (Leinco C2268) or control IgG (Leinco R1367) antibody were delivered i.p. beginning on Day 8, and MPLA + CoA treatments initiated on Day 11. Supplemental antibody injections (500ug) were given every 4-5 days through the course of the experiment to maintain TAM depletion, which was confirmed by flow cytometry. For checkpoint blockade experiment, mice receiving all treatments, including 10mg/kg i.p. anti-CTLA4 checkpoint blockade therapy (clone 9D9, Platinum grade, Leinco), were treated a total of 6 times between Day 11 and Day 25. Treatments were then discontinued and survival monitored. Animals were sacrificed when tumors reached >20mm in any dimension.

### Flow cytometry

For TAM flow cytometry, 50mg of freshly excised tumor tissue was transferred to a 1.5mL tube, manually dissociated with scissors, and digested in DMEM with 0.1% collagenase (Sigma C6885) for 1.5 hour at room temperature with gentle shaking. Sample was passed through a 100μm filter, RBCs lysed, and cell pellet stained on ice in FACs buffer (PBS with 10mM HEPES and 5% BSA). After Fc Block (1:100, BD Biosciences 553142), cells were labeled with anti-CD45-PeCy7 (clone 30-F11, BD Biosciences 552848, 1:400), anti-F4/80-PE (clone BM8, eBioscience 12-4801-82, 1:200), and anti-CD11b-APC (clone M1/70, BioLegend 101225, 1:200). Cells were washed and resuspended in FACs buffer supplemented with DAPI to exclude dead cells, and live CD45+ F4/80+ CD11b+ TAMs quantified using an LSR II analyzer (BD Bioscience) running BD FACSDiva software. Data were analyzed with FlowJo (Tree Star, v10.8.1). For BMDM viability assessment after CoA supplementation, media was removed and cells washed once with PBS. Cells were then incubated in PBS with 4% EDTA supplemented with DAPI and fluorescein diacetate (FDA, Thermo Fisher F1303) at 37 °C for 5 min. Cells were then lifted by gentle pipetting, and the percentage of DAPI negative, FDA positive live cells per total events per well quantified using the aforementioned cytometer and software.

For H3K27ac intracellular staining-flow cytometry, cells were treated as indicated, washed and Fc-blocked (BD 553141, 1:500) on ice, and processed with BD Cytofix/Cytoperm Fixation and Permeabilization Kit (BD 554714). Anti-H3K27ac primary antibody (Sigma SAB4200485) was used at 1:500, and secondary anti-rabbit 488 (Invitrogen) at 1:1000. Trichostatin A (TSA) treatment served as positive control.

For cell surface staining of macrophage activation markers, following Fc-blocking, cells were stained on ice with the following antibodies (all 1:500): rat isotype PE (BioLegend 400507), CD80 PE (eBioscience 12-0801-82), CD86 PE (BioLegend 105008), H-2Kd PE (BioLegend 116607), IFNGR PE (eBioscience 12-1191-82), CD64 APC (BioLegend 139306), MHC Class II APC780 (eBioscience 47-5321-82).

### Intratumoral cytokines ELISA

Approximately 30mg of flash frozen, mortar and pestal-homogenized PyMT tumor tissue was transferred to an 1.5mL tube and incubated in 350uL of PBS at 37°C with shaking for 45 minutes to wash and collect intratumoral cytokines. Tumor tissue was then pelleted by centrifugation, and cytokine levels in the supernatant/PBS wash quantified using ELISA kits according to manufacturer’s instructions (murine IL-1β: Invitrogen 88-7013-22, murine IL-6: Invitrogen 88-7064-88) and a SpectraMax M2 plate reader. A small number of suspected outliers were removed from some data sets using the ROUT method, leading to some discrepancies between the experimental “n” in tumor growth plots versus “n” presented in the ELISA results.

### *L. pneumophila* bacterial strains and *in vitro* BMDM infection

*Legionella pneumophila* Δ*flaA*, derived from the virulent clinical isolate JR32 (*89*), was engineered to express the *Photorhabdus luminescens luxCDABE* operon as previously described (*90*). Bacteria from frozen glycerol stocks were cultured first on buffered yeast extract with charcoal (BCYE) agar plates, then in buffered yeast extract (BYE) broth supplemented with L-cysteine, ferric nitrate, and streptomycin (100ug/mL). For infection, BMDMs differentiated in RPMI/3T3 supernatant as described above were plated at 100K per well in 96-well opaque white plates in RPMI without antibiotics. The following day, BMDMs were infected with bacteria from an overnight liquid culture diluted in antibiotic-free RPMI for 1 hour at 37 °C at a MOI of 0.01. After replacing the infection media with RPMI containing H_2_O vehicle, CoA, or IFNγ (6 ng/mL), a Day 0 luminescence measurement was recorded (λ470, Spectramax L luminometer, Molecular Devices). For the 5mM CoA condition, sterile NaOH was used to correct the pH of the media, which was acidified by this high CoA concentration. Intracellular bacterial growth was then tracked over time by reading luminescence every 24 hours, with fresh CoA supplemented into the appropriate wells after each reading. At the experimental endpoint, BMDMs were lysed with water and lysate dilutions were plated on BYCE agar plates to enumerate CFU.

### Metabolomics

For metabolomics in Figure 2B, BMDMs were cultured in DMEM (Corning 17-207-CV) supplemented with 10% One Shot dialyzed FBS (Thermo Fisher A33820-01), 1mM sodium pyruvate (Gibco 11360-070), 4mM L-glutamine (Gibco 25030081), and 25mM glucose (Gibco A24940-01). 5×10^5^ cells were first pretreated with 250µM CoA or acetyl-CoA for 2h hours, followed by EtOH or 5µM 4-OHE1 for 1 hour. Cells were then stimulated with 100ng/mL LPS for 1.5 hours. For the last 30 minutes post-LPS stimulation, cells were washed with unsupplemented media, and provided DMEM described above except with 25mM ^13^C_6_-glucose (Cambridge Isotope Laboratories, CLM-1396-PK) to competitively label metabolites between conditions as a proxy for metabolic pathway activity. After this 30 minutes of labeling, cells were washed with PBS and moved to dry ice for addition of 1mL ice cold 80% MeOH. Cells were incubated at −80°C for 15 minutes, scraped on dry ice, and moved to low bind Safe-Lock 1.5mL tubes (Eppendorf cat.022363204). After centrifugation 20,000g x 10 min at 4°C, the cell pellet was saved for protein quantification by BCA assay, while the MeOH extract was transferred to a new tube and evaporated overnight at RT using a SpeedVac. Dried extracts were stored at −80°C. For metabolomics experiments in Figures S2C and S2D, we utilized DMEM with a lower glucose concentration (10mM, in line with RPMI glucose concentrations used for similar tracing by other groups (*22*)). This allowed better detection of LPS-induced ^13^C6-glucose labeling of TCA cycle metabolites, potentially via reduction of basal glycolytic spillover, and/or reduction of the Crabtree effect. The media was prepared by supplementing glucose-free DMEM (Gibco 11966-25) with 10mM glucose (Gibco A24940-01) and 0.5 mM sodium pyruvate (Gibco 11360-070).

For whole tissue metabolomics, approximately 30mg of flash frozen, mortar-and-pestal ground tumor tissue was transferred to a low bind 1.5mL tube. 500 microliters of 80% MeOH was added to the tube, and tissue further homogenized with handheld electrical tissue grinding probe. Following rigorous vortexing and two freeze/thaw cycles in the −80°C freezer, samples were centrifuged 20,000g x 10 min at 4°C. The cell pellet was saved for protein quantification by BCA assay, while the MeOH extract was transferred to a new tube and evaporated at RT using a SpeedVac.

Dried MeOH extracts were resuspended in 50% ACN:water and 1/10^th^ was loaded onto a Luna 3um NH2 100A (150 × 2.0 mm) column (Phenomenex). The chromatographic separation was performed on a Vanquish Flex (Thermo Scientific) with mobile phases A (5 mM NH4AcO pH 9.9) and B (ACN) and a flow rate of 200µL/min. A linear gradient from 15% A to 95% A over 18 min was followed by 9 min isocratic flow at 95% A and reequilibration to 15% A. Metabolites were detected with a Thermo Scientific Q Exactive mass spectrometer run with polarity switching (+3.5 kV/− 3.5 kV) in full scan mode with an m/z range of 65-975. TraceFinder 4.1 (Thermo Scientific) or Maven v8.1.27.11 was used to quantify the targeted metabolites by area under the curve using expected retention time and accurate mass measurements (< 5 ppm). Values were normalized to sample protein concentration. Relative amounts of metabolites were calculated by summing up the values for all isotopologues of a given metabolite. Fraction contributions (FC) of ^13^C carbons to total carbon for each metabolite was calculated as previously described (*100*). Data analysis was performed using in-house R scripts available upon request.

### H3K27ac Chromatin Immunoprecipitation (ChIP) qPCR

ChIP was performed as previously described (*34*). One plate of supBMDMs (approximately 10 millions cells) was used per ChIP. 5μL of anti-H3K27ac antibody (Cell Signaling 8173) was used per H3K27ac ChIP, with an equal μg amount of rabbit IgG used in control ChIPs. For each individual H3K27ac or IgG control ChIP, inputs were diluted and used to generate standard curves to calculate the percent enrichment relative to input for each sample for each genomic fragment analyzed. Primers used are as follows:

*Il1b 2kb enhancer -* GCCTGAGTGTCAGCTCCAAG, GCCTGAGTGTCAGCTCCAAG

*Il1b 10kb enhancer –* TGCTCTGACCACTTACTGCC, GAAAACGGTCCAGCAATGGG

*Il1b control region -* GCAAACTCCGGTGCGAGTA, GCATCTGAGGCACCTACTGAA

*Il1a promoter -* TATTGTGCCCCAGGACTGAA, CCTCAAATGCCTTAGCTGCC

*Il1a 10kb enhancer-* AAGGACAGAGCTGGAGGGAG, TGAGAGGTCTTTCCACCAAGG

*Il1a control-* GCCCAGAAGAAGATGCAAACTC, CCCTACAATAAAATAGCAGCTGACC

*Nr1h2 promoter –* TGTCCGCACCGGTGATTG, ACTCCCAGGCTTCTGAAGTTAC

*Cox5a promoter –* CGTCTGGAGTCGTGTAGGAG, GCAAGTCCACACACCTGACT

*Nop14 promoter –* TTAGTTTTCGCCGGGCCTC, GCACTGTCGTCCAAAGTGTG

### H3K27ac Cut-and-Tag

H3K27ac Cut-and-Tag was performed as described (*101*) using in-house-prepared pA-MNAse, with 2-3 independent biological replicates per experimental condition. 500,000 rBMDMs per condition were pretreated with vehicle or CoA (500μM) for 2h before 30min vehicle or LPS stimulation (100ng/mL). H3K27ac antibody (Sigma SAB4200485) was used at 1:50. Tagmented DNA was purified with Zymo ChIP DNA Clean & Concentrator columns. DNA was amplified with Phusion polymerase using previously described primers (*102*) for multiplexing, pooling, and sequencing on an Illumina NovaSeq 6000 (50bp PE reads). Data were aligned to mm10 with RNASTAR (*103*) and peaks called and quantified with HOMER (*16*).

### 2-NDBG uptake

The fluorescent glucose analog 2-NDBG (Invitrogen N13195) was added directly to cell culture media at a final concentration of 5mg/ml. Cells were then washed with PBS on ice, lifted with PBS/5mM EDTA with DAPI, and fluorescence analyzed on a BD LSR II flow cytometer.

### Endotoxin testing

Pierce Chromogenic Endotoxin Quant Kit (Thermo Scientific A39552) was used to asses endotoxin levels in 10mM solutions of all CoA/CoA precursors/CoA degradation products used in this study. All were reconstituted in the same water (Invitrogen UltraPure DNAse/RNAse Free water, 10977015), which meets USP WFI (water for injection) standards of <0.25 endotoxin units per mL. To estimate endotoxin exposure in cells treated with 500μM of any of the assayed CoAs, CoA precursors, and CoA degradation products, we used the following conversion: 10 EU/mL = 1.0 ng/mL of endotoxin/LPS.

### CoA tracing

For quantification and isotope tracing, Acyl-CoAs were measured by liquid chromatography-high resolution mass spectrometry as previously described in detail (*104*). Briefly, media was aspirated, and cells were quenched in 1 mL 10% (w/v) trichloroacetic acid in water, scraped, and extracted. Samples were homogenized by using a probe tip sonicator in 0.5 second pulses 30 times then centrifuged at 17,000 x g for 10 min at 4°C. Samples were purified by solid phase extraction (SPE) cartridges (Oasis HLB 10 mg, Waters) that were conditioned with 1 mL of methanol and 1 mL of water. Acid-extracted supernatants were loaded onto the cartridges and washed with 1 mL of water. Acyl-CoAs were eluted with 1 mL of 25 mM ammonium acetate in methanol and evaporated to dryness under nitrogen. Samples were resuspended in 50 µL of 5% (w/v) 5-sulfosalicyilic acid and 20 µL injections were analyzed on an Ultimate 3000 UHPLC using a Waters HSS T3 2.1×100mm 3.5 µm column coupled to a Q Exactive Plus mass spectrometer. The analysts were blinded to sample identity during processing and quantification. Data was analyzed using Tracefinder 5.1 (Thermo Scientific).

### NFkB luciferase

RAW and THP-1 cell lines with stably integrated NFkB luciferase reporter constrict were a gift from Dr. Gregory Barton (UC Berkeley). Following treatments, cells were washed 5 times in PBS to get rid of extracellular CoA (which positively influences Firefly Luciferase activity as it is a component of the lysis/assay buffer). Cells were then lysed in 100 microliters of 1x Firefly Assay Buffer (0.15M Tris pH 7.5, 75mM NaCl, 3mM MgCl_2_, 0.25% Triton-100, 5mM DTT, 0.2mM CoA, 0.15mM ATP, and 1.4mg/mL luciferin). After 10 minutes, lysates were moved to a white bottom plate and luciferase signal read on a Spectramax M2 microplate reader.

### Statistical analysis

Majority of statistical analysis was performed using GraphPad Prism 9 software. Metabolomics statistical analysis was performed using R or GraphPad Prism 9. All data were assumed to be normally distributed. One-Way ANOVA was used in initial metabolomics FC analysis comparing multiple groups, with Dunnett’s post-test used to identify significant differences between specific groups. In other experiments, planned comparisons were built-in to experimental design as indicated, with Student’s t-test used to make sensible comparisons between two experimental groups. Metabolomics data were log-transformed and Z-scored for representation on heatmaps. Where indicated in the methods (see above), suspected outliers were removed from experimental groups using the ROUT method.

### Illustrations

Images were created using BioRender and Adobe Illustrator (licenses to G.A.T.).

---

**Figure S1.**
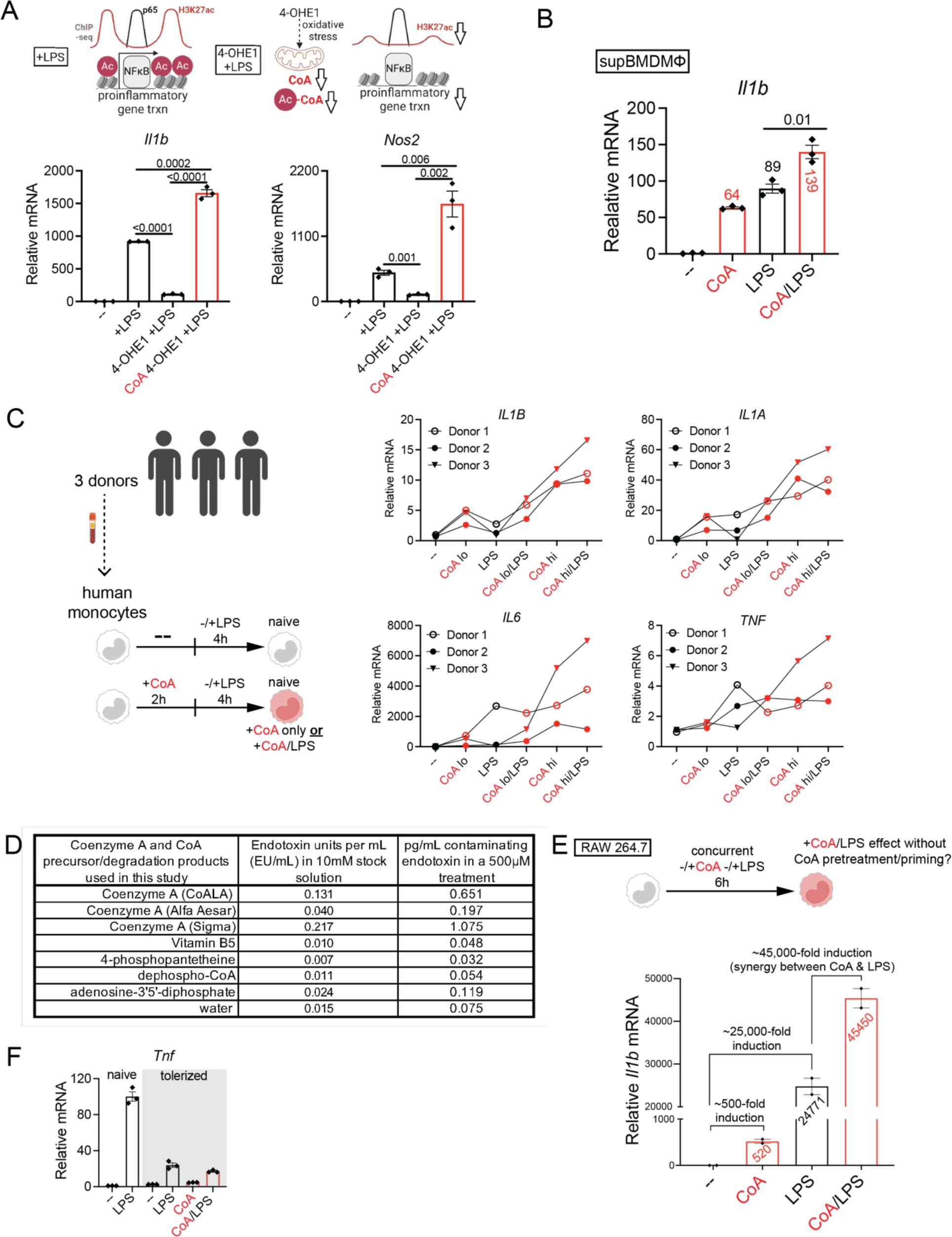
**A.** Top – Illustrated summary of 4-OHE1’s disruption of CoA and acetyl-CoA homeostasis and LPS-induced histone acetylation. Bottom - qPCR for LPS-induced *Il1b* and *Nos2* in BMDMs (1.5h) pretreated for 1h with 4-hydroxyestrone (5μM) and/or CoA (250μM). **B.** qPCR for LPS-induced *Il1b* (2h) with CoA supplementation (250μM) in naïve BMDMs differentiated with MCSF-containing supernatant from 3T3-L1 cells (supBMDMs). **C.** qPCR for proinflammatory gene expression in primary human monocytes from 3 independent donors with CoA supplementation at two doses (lo = 500μM CoA, hi = 5mM CoA) either alone or with LPS. **D.** Endotoxin levels in CoAs and CoA precursors/degradation products utilized in this study, and estimated levels of contaminating endotoxin in a 500μM *in vitro* CoA treatment. **E.** qPCR for *Il1b* in RAW264.7 macrophages treated with CoA (250μM), LPS, or simultaneously cotreated with CoA/LPS, for 6h. **F.** qPCR for LPS-induced *Tnf* expression in naïve or tolerized rBMDMs with CoA supplementation (2.5mM). For qPCRs in **A**, **B**, **E**, and **F**, n=2 or 3 biological replicates per condition, mean ± s.e.m is shown, numbers above bars are Student’s t-test P values (planned comparisons).

**Figure S2.**
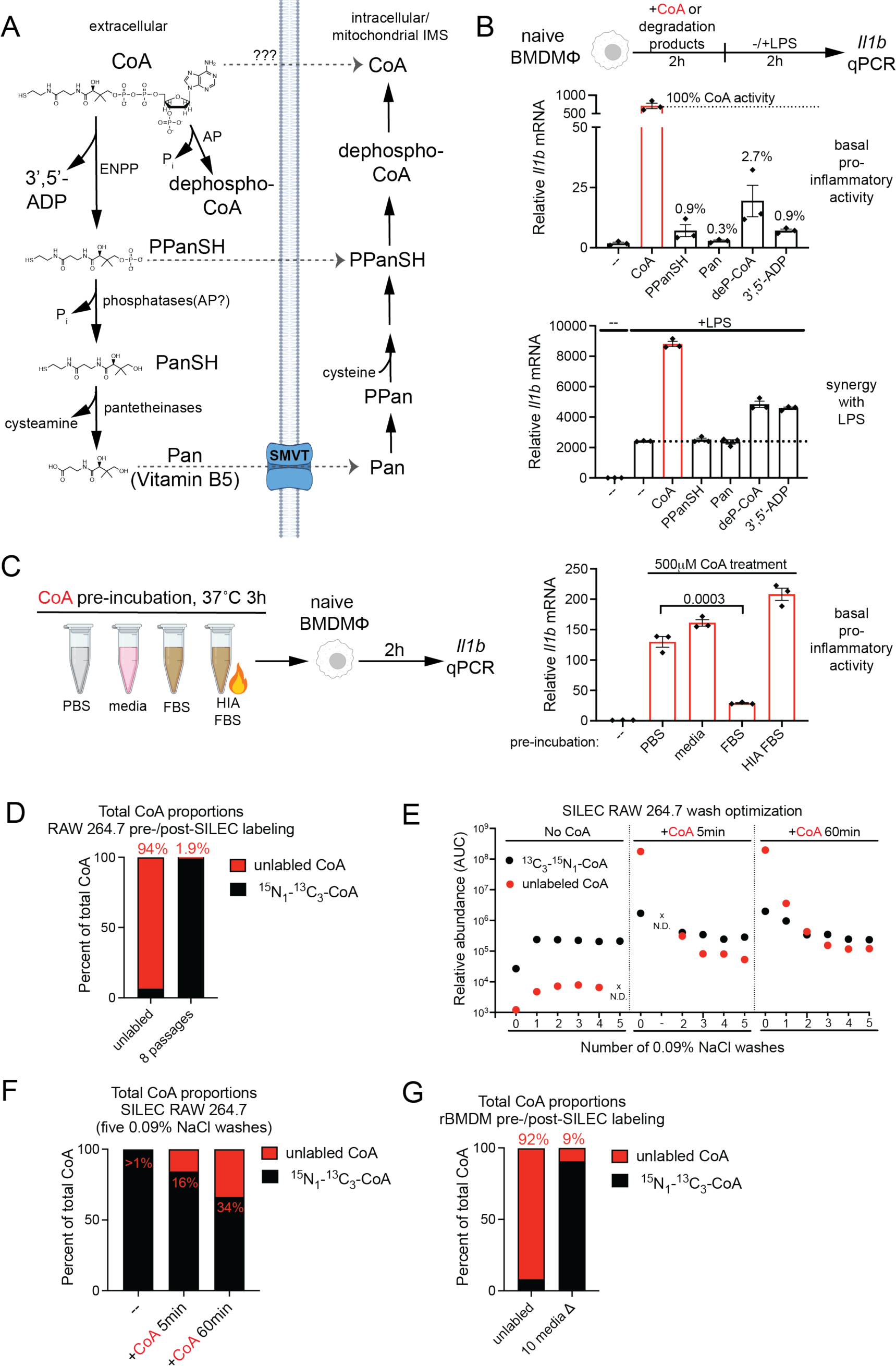
**A.** Schematic depicting known pathways of extracellular CoA degradation product entry into intracellular CoA biosynthesis pathway. **B.** *Il1b* qPCR in rBMDMs treated with CoA or CoA degradation products (500μM). Percentages represent *Il1b* induction relative to CoA. **C.** *Il1b* qPCR in rBMDMs treated with CoA (500μM) for 2h. Prior to addition to rBMDMs, CoA was preincubated at 37°C for 3h in PBS, media, non-heat inactivated FBS, of heat-inactivated (HIA) FBS. For qPCRs in **B** and **C**, n=4, and mean ± s.e.m. is shown, and numbers above data are Student’s t-test P values. **D**. Unlabeled CoA as a percentage of the total CoA pool (set to 100%) pre- and post-SILEC labeling of RAW264.7 cells. n=1. **E**. Unlabeled and labeled CoA levels in SILEC RAW264.7 cells treated with unlabeled CoA for 5 or 60 minutes and subjected to the indicated numbers of 0.9% NaCl washes. n=1, mean is shown. **F**. Unlabeled CoA as a percentage of the total CoA pool (set to 100%) in SILEC RAW264.7 cells treated with unlabeled CoA for the indicated times and subjected to 5 0.9% NaCl washes. n=1. **G**. Unlabeled CoA as a percentage of the total CoA pool (set to 100%) pre- and post-SILEC labeling of rBMDMs. n=1.

**Figure S3.**
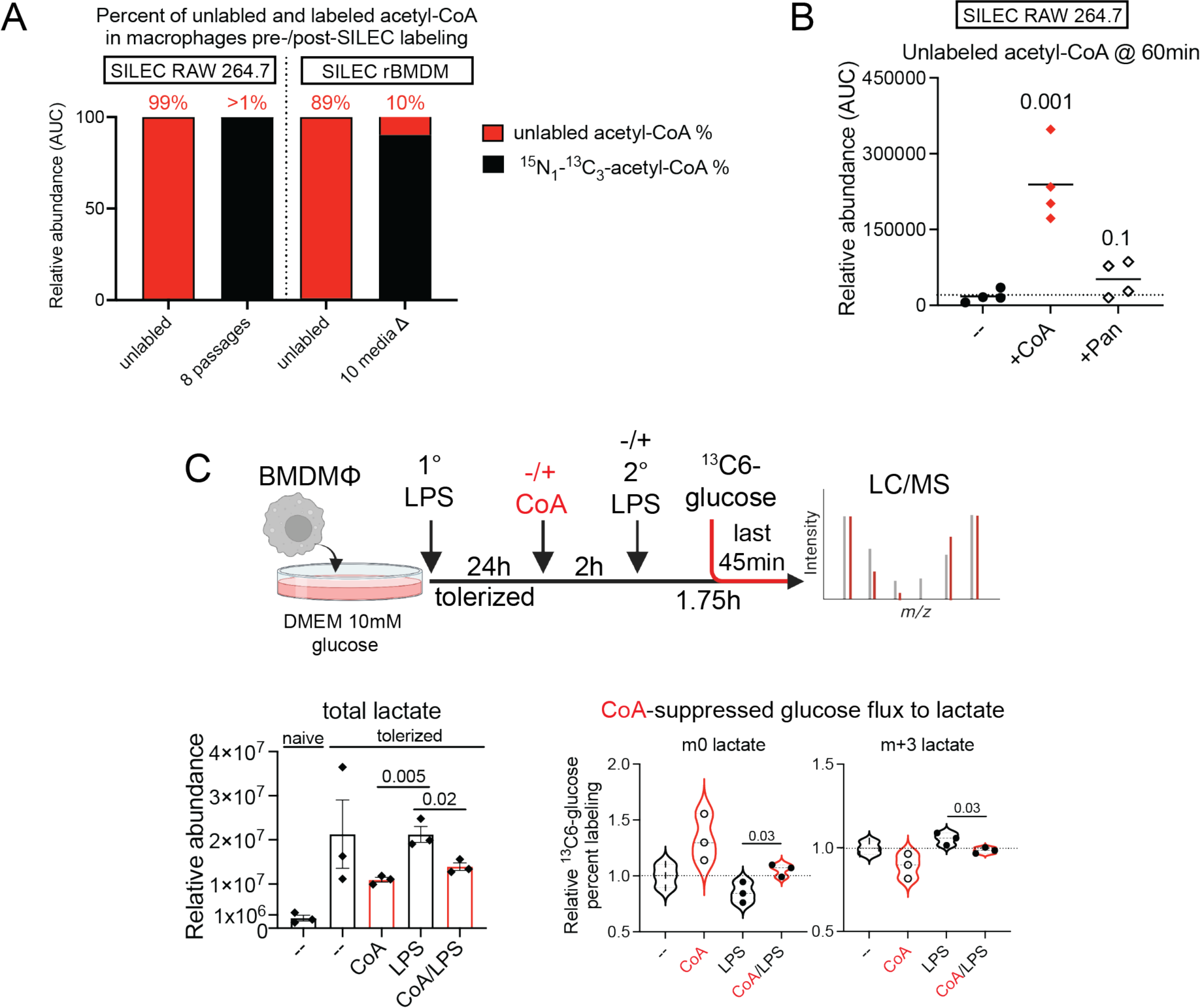
**A**. Unlabeled acetyl-CoA as a percentage of the total CoA pool (set to 100%) pre- and post-SILEC labeling of RAW264.7 cells (left) and rBMDMs (right). n=4. **B**. Unlabeled acetyl-CoA levels in SILEC RAW264.7 macrophages 60 minutes following unlabeled CoA or Pan treatment (both 500μM). n=4, mean is shown, numbers above data are Student’s t-test P values (planned comparison with the untreated condition). **C**. Total lactate levels (left), and relative ^13^C_6_-glucose fractional contribution (FC) to lactate (right), in LPS-tolerized BMDMs treated with CoA (2.5mM), LPS, or CoA/LPS. n=3 biological replicates per condition, mean ± s.e.m is shown, numbers above bars are Student’s t-test P values (planned comparisons).

**Figure S4.**
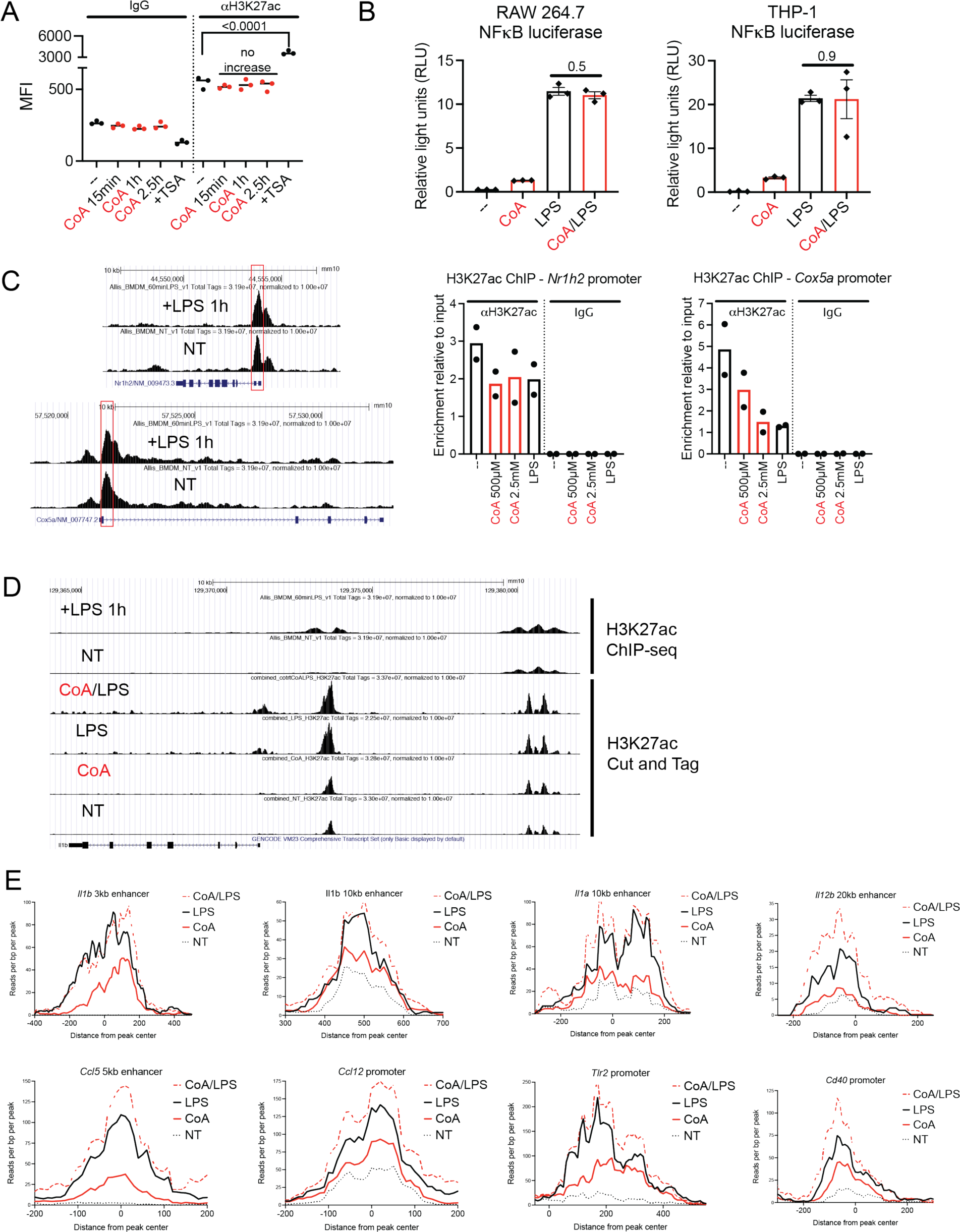
**A**. Quantification of Fig.4B H3K27ac ICS-flow cytometry experiment assessing global H3K27ac upon CoA treatment. n=3, mean is shown. **B**. Luciferase activity in RAW264.7 (left) and THP-1 cells (right) with stably-integrated NFκB luciferase reporters following 2 hour treatment with either CoA (500μM), LPS, or CoA/LPS co-treatment. n=3, mean ± s.e.m. is shown, and numbers above bars are Student’s t-test P values (planned comparisons). **C**. H3K27ac levels at *Nrh1h2* and *Cox5a* promoters assessed by ChIP-qPCR in cells treated with CoA or LPS. **D**. Comparison of *Il1b* locus H3K27ac peaks captured in previously published ChIP-seq experiment from reference 49, and by our H3K27ac CnT experiments. **E**. CnT read density at putative *cis*-regulatory region H3K27ac peaks in the indicated genes in rBMDMs treated with either CoA (500μM), LPS, or CoA/LPS co-treatment.

**Figure S5.**
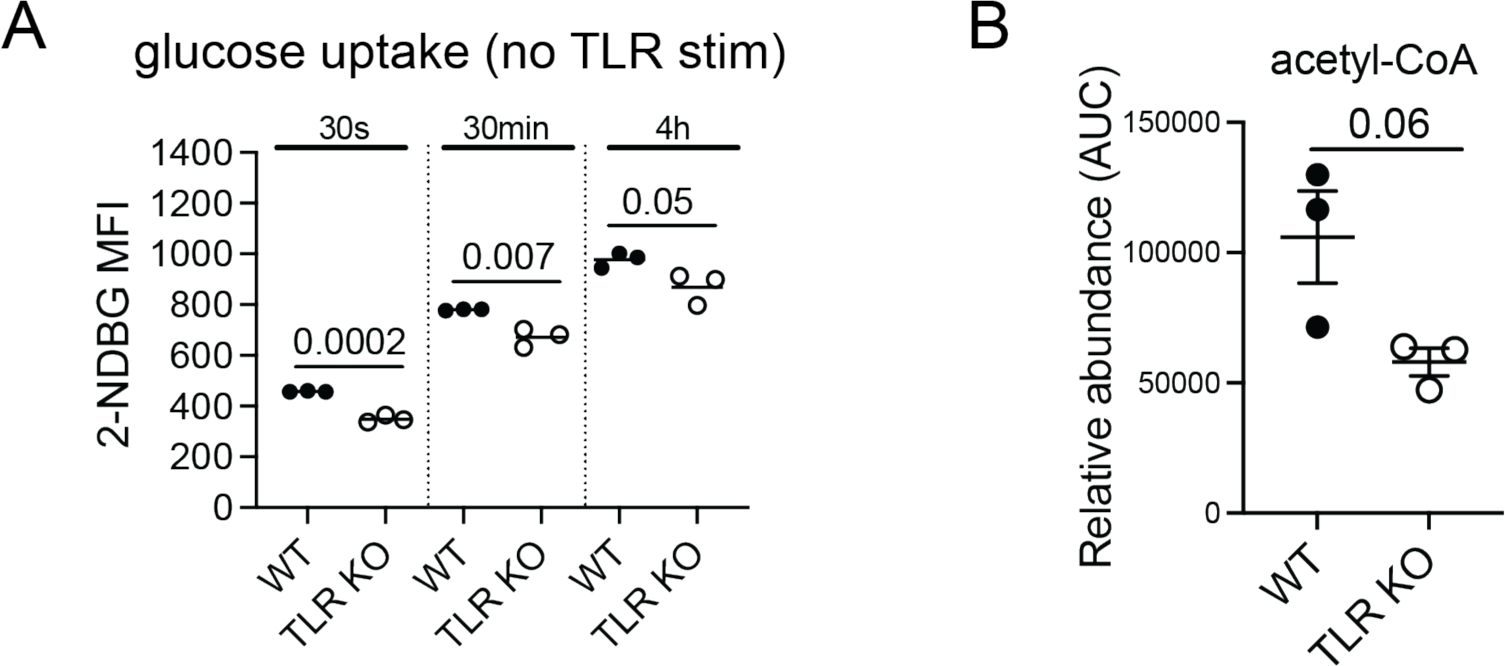
**A**. 2-NDBG uptake in WT vs. TLR KO supBMDMs. n=3, mean is shown, numbers above bars are Student’s t-test P values (planned comparisons). **B**. Steady-state levels of acetyl-CoA in WT versus TLR KO rBMDMs measured as part of Fig.5C experiment.

**Figure S6.**
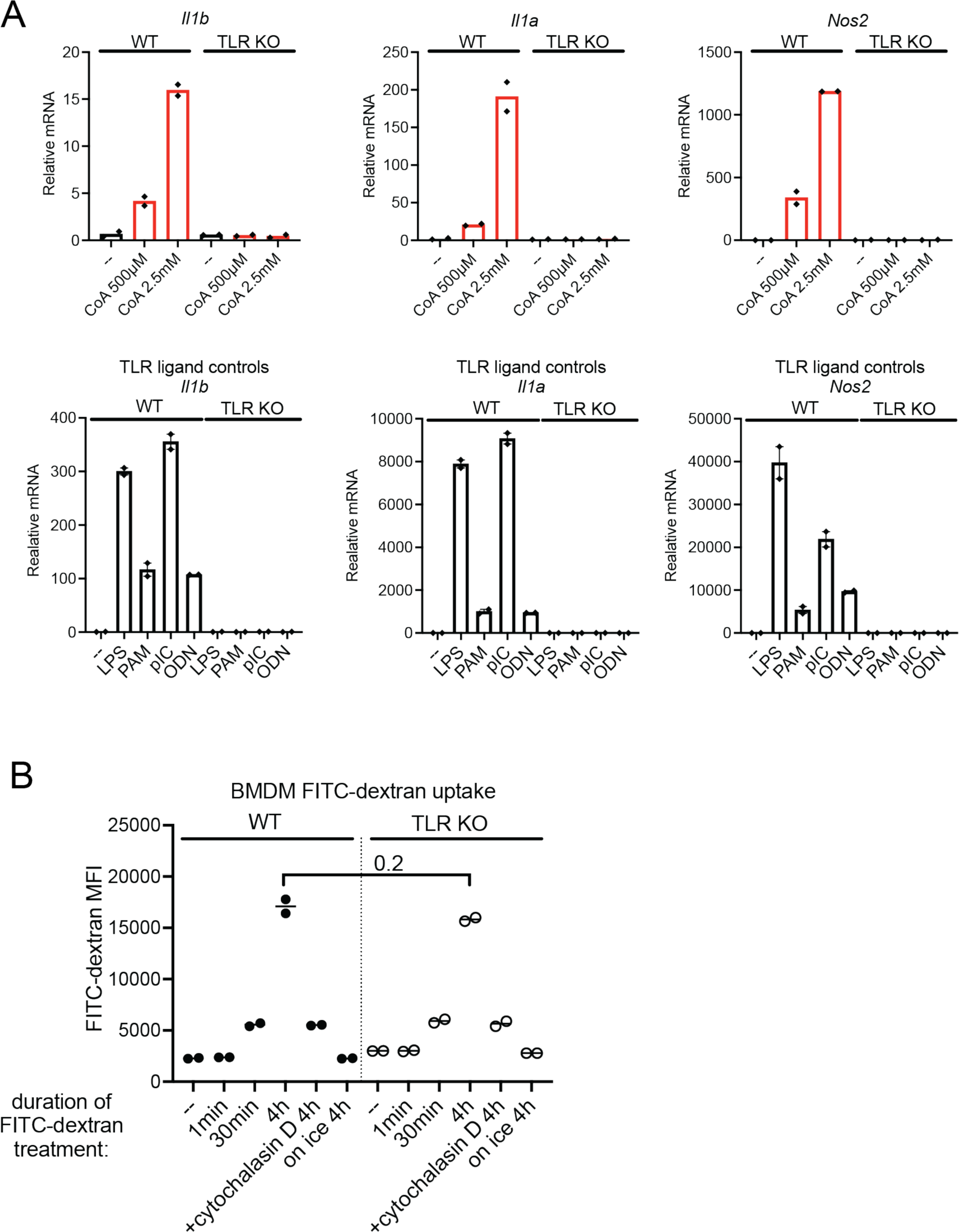
**A**. *Il1b*, *Il1a*, and *Nos2* expression measured by qPCR in rBMDMs treated with CoA (top), or TLR ligands (bottom). n=2, mean is shown. **B**. Flow cytometry measurement of FITC-Dextran uptake over time in WT versus TLR KO rBMDMs. n=2, mean is shown.

**Figure S7.**
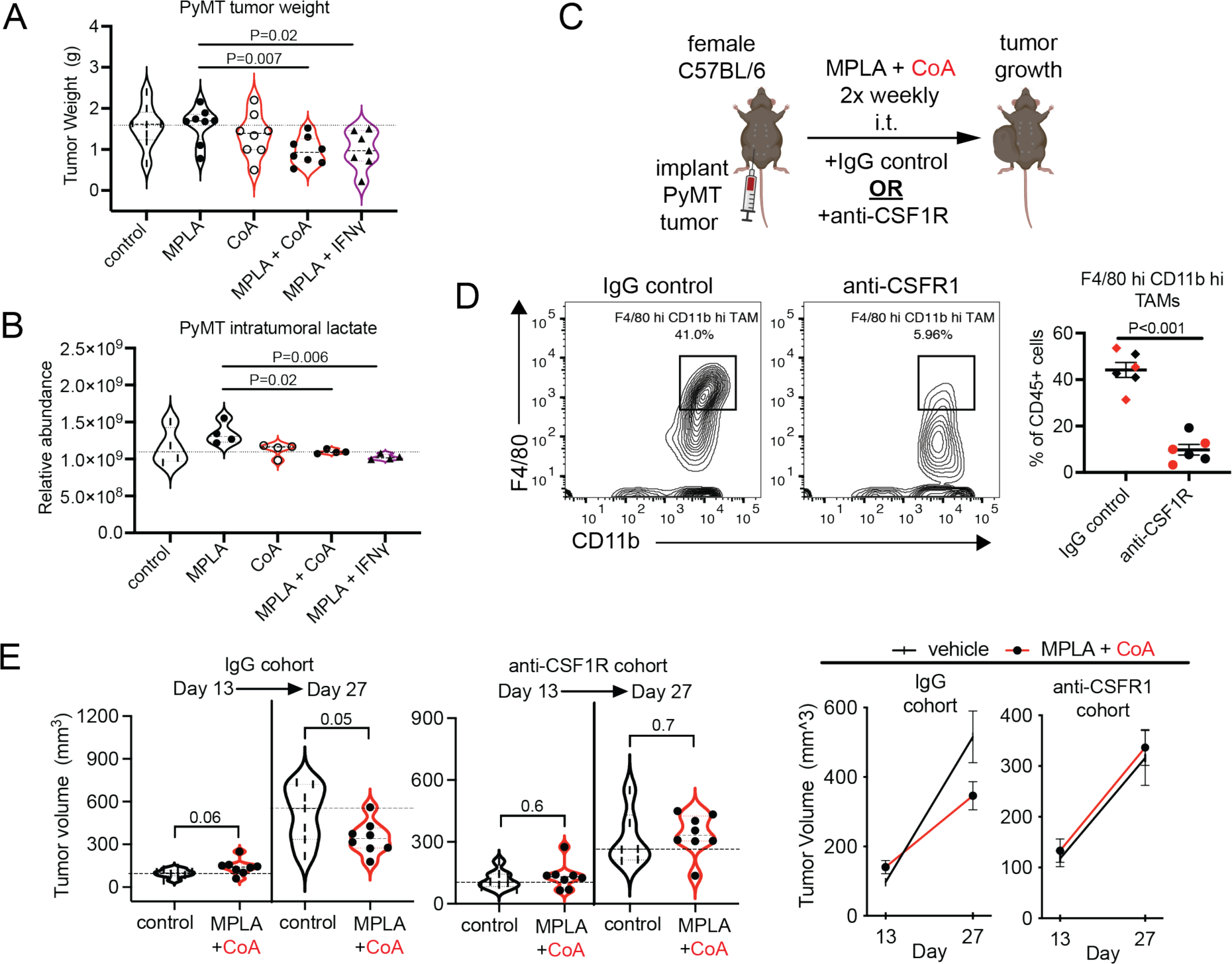
**A, B.** Day 30 tumor weights (A) and intratumoral lactate level (B) for tumors from mice in Fig. 7B experiment. **C.** Top - Experimental strategy to test the contribution of tumor-associated macrophages (TAMs) to the anti-tumor effect of MPLA+CoA therapy. Bottom - Effects of TAM depletion on MPLA+CoA therapy efficacy, represented by comparing tumor size among all animals (left), or by comparing tumor growth rates (right), on experimental Days 13 and 27 between IgG and anti-CSFR1 cohorts receiving vehicle or MPLA+CoA therapy. n=8 or 7 mice per group. **D.** Left – flow cytometry of CD45+ tumor cells depicting F4/80 hi CD11b hi TAM populations in representative mice from IgG control antibody-treated or anti-CSF1R antibody-treated cohorts. Right – quantification of flow cytometry results confirming TAM depletion in anti-CSF1R-treated mice. 6 mice were chosen at random from the IgG control or anti-CSF1R cohorts. Black indicates mice receiving vehicle control treatment, red indicates mice receiving MPLA+CoA therapy. All numbers above bars in figure represent Student’s t-test P values (planned comparisons).

**Figure S8.**
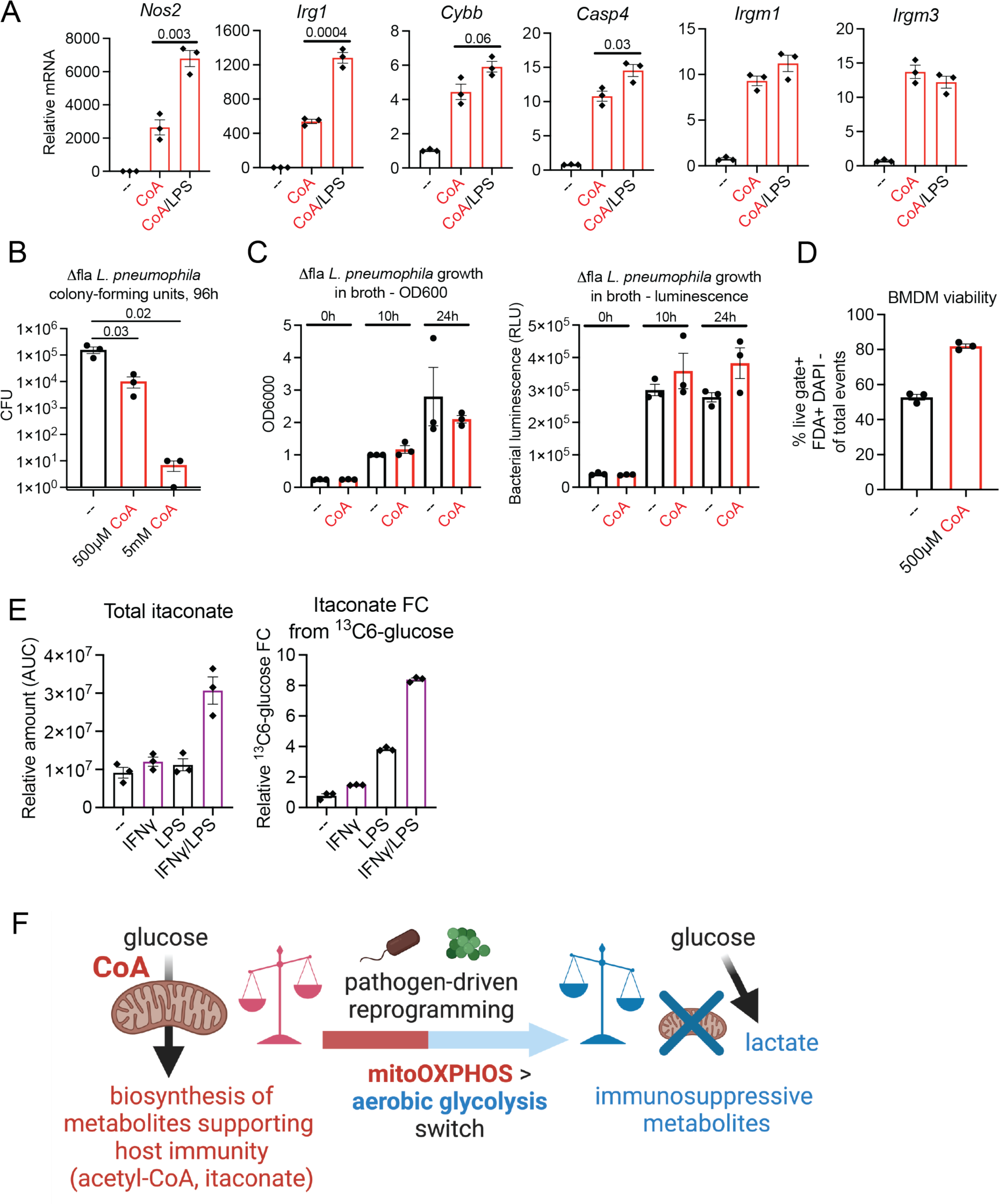
**A.** qPCR for *L. pneumophila* restriction genes in rBMDMs treated with CoA alone (250μM), or CoA in combination with LPS, for 2h. n=3, mean ± s.e.m is shown, numbers represent Student’s t-test P values (planned comparisons). **B.** *L. pneumophila* colony forming units (CFUs) from infected BMDMs in Fig. 8A 96h after infection. n=3, mean ± s.e.m is shown, numbers represent Student’s t-test P values (planned comparisons). **C.** *L. pneumophila* growth (left) and luminescence (right) in broth supplemented with 500μM CoA for 24 hours. n=3, mean ± s.e.m is shown. **D.** supBMDM viability assessed by flow cytometry after 72h of continuous 500μM CoA supplementation. n=3, mean ± s.e.m is shown. **E.** Total steady-state itaconate levels (left), and relative ^13^C_6_-glucose fractional contribution (FC) to itaconate (right), in rBMDMs treated with IFNγ (xxx), LPS, or IFNγ/LPS for 2h. **F.** Model depicting how a pathogen-driven mitoOXPHOS to aerobic glycolysis shift reduces production of proinflammatory/anti-microbial metabolites to create a replicative niche for pathogens.

## Notes

### Competing Interest Statement

Greg A. Timblin and Joshua N. Farahzad are co-founders and own equity in Inapill Inc.

### Summary of Updates

In this updated version of the manuscript, we: 1) collaboratively developed a novel isotopic labeling system to allow CoA pulse-chase studies. This has allowed acquisition of data supporting the main concept of the study: that macrophages are indeed able to take up exogenous CoA to support proinflammatory immunometabolism. This technical advance provides key mechanistic insight into the anti-tumor and anti-microbial effects of CoA via enhancement of specific CoA-supported aspects of host metabolism. It is also, to our knowledge, the first demonstration using a Vitamin B5-based tracing strategy that this key metabolic co-factor can be directly acquired by mammalian cells (or any eukaryotic cell type). 2) utilized both classic (ChIP) and current (Cut-and-Tag) technologies to provide new, extensive characterization of the epigenetic changes that the metabolic effects of CoA have on macrophage histone acetylation. These targeted epigenetic changes, which we show occur not globally, but specifically at proinflammatory gene loci, provide a mechanistic basis for the anti-tumor and anti-microbial phenotype that CoA instills in macrophages. 3) with regards to the promotion of an anti-microbial macrophage phenotype by CoA, we genetically link this effect to the Acod1/Irg1 gene and the production of itaconate. Convincingly, we show that CoA not only promotes the expression of IRG1, but also that CoA augments the biosynthesis of glucose-derived itaconate through its ability to support and drive mitochondrial glucose oxidation via the TCA cycle. 4) we provide strong evidence that the proinflammatory effects of CoA rely on tonic TLR signaling, which we thoroughly demonstrate is a critical regulator of the basal metabolic, epigenetic, and phenotypic state of macrophages. This novel concept, which has broad implications for immunologist (particularly those studying TLR biology), could only be investigated using our murine TLR KO system.

